# Exogenous estradiol and oxytocin modulate sex differences in hippocampal reactivity and episodic memory

**DOI:** 10.1101/2021.11.22.469500

**Authors:** Marie Coenjaerts, Isabelle Trimborn, Berina Adrovic, Birgit Stoffel-Wagner, Larry Cahill, Alexandra Philipsen, René Hurlemann, Dirk Scheele

**Affiliations:** Division of Medical Psychology, Department of Psychiatry and Psychotherapy, University Hospital Bonn, 53105 Bonn, Germany; Institute for Clinical Chemistry and Clinical Pharmacology, University of Bonn, 53105 Bonn, Germany; Department of Neurobiology and Behavior, University of California, Irvine, CA, 92697-3800, USA; Department of Psychiatry and Psychotherapy, University Hospital Bonn, 53105 Bonn, Germany; Department of Psychiatry, School of Medicine & Health Sciences, University of Oldenburg, 26129 Oldenburg; Research Center Neurosensory Science, University of Oldenburg, 26129 Oldenburg

**Author notes:** **Corresponding authors:** Marie Coenjaerts, Department of Psychiatry and Psychotherapy University Hospital Bonn, Venusberg-Campus 1, 53127 Bonn, Germany. **Corresponding authors:** Dirk Scheele, Department of Psychiatry and Psychotherapy University Hospital Bonn, Venusberg-Campus 1, 53127 Bonn, Germany.

**Keywords:** episodic memory, estradiol, hippocampus, oxytocin, sex differences

## Abstract

Considerable evidence supports sex differences in episodic memory, which may translate to heightened vulnerability to stress- and trauma-related disorders in women. The hormones estradiol and oxytocin both affect episodic memory, but possible underlying hormonal interactions have not been systemically tested in humans. To this end, healthy women (n = 111) and men (n = 115) received estradiol gel (2 mg) or placebo before the administration of intranasal oxytocin (24 IU) or placebo in a randomized, placebo-controlled, parallel-group functional magnetic resonance imaging (fMRI) study. In the fMRI session, participants viewed positive, neutral, and negative scenes. A surprise recognition task was conducted three days later. Under placebo, women showed a significantly better recognition memory and increased hippocampal responses to subsequently remembered items independent of the emotional valence compared to men. The separate treatments with either hormone significantly diminished this mnemonic sex difference and reversed the hippocampal activation pattern. However, the combined treatments led to a memory performance comparable to that of the placebo group. Collectively, the results suggest that both hormones play a crucial role in modulating sex differences in episodic memory. Furthermore, possible antagonistic interactions between estradiol and oxytocin could explain previously observed opposing hormonal effects in women and men.

## Introduction

Sex differences in memory have been reported repeatedly^[1–3]^ and may contribute to the higher prevalence of stress- and trauma-related disorders in women.^[4]^ Women tend to outperform men in episodic memory functions,^[1,3,5,6]^ including autobiographical and recognition memory,^[7,8]^ while there appears to be an advantage for spatial-based memory tasks in men.^[5]^ Moreover, compared with men, women show enhanced memory for emotional stimuli.^[9]^ Emotional hypermnesia is partially mediated through activation of the amygdala^[10,11]^ and imaging studies revealed a sex-related hemispheric lateralization of the amygdala in response to emotional stimuli: right amygdala activation while viewing emotional stimuli is more significantly related to subsequent memory for the images in men than women, whereas the reverse sex difference is evident for the left amygdala.^[9,12–14]^

Potential biological mechanisms explaining these sex differences include the effects of sex hormones on memory function and neuroplasticity.^[1,15]^ In particular the role of estradiol (EST), as the most potent and prevalent circulating estrogen is discussed. As EST elicits effects on the memory consolidation in fear extinction learning, a behavioral process that models the psychopathology of PTSD and anxiety disorders,^[4]^ the regulating function of mnemonic brain regions, such as the hippocampus, has been highlighted.^[16]^ Previous studies in rodents and humans have shown that hippocampal function is sensitive to changes in estrogens that occur across the reproductive cycle, pregnancy, or ageing.^[15,17,18]^ EST affects the dendritic spine density in the hippocampus^[3,16]^ and enhances neuronal growth by promoting the formation of new synaptic connections.^[19]^ Several rodent studies have shown that the infusion of EST before or immediately after a training improves memory performance.^[16,20–24]^ Correlative studies of natural hormonal fluctuations during the menstrual cycle found that estrogen levels are associated with greater episodic memory performance, but estrogen exposure has not been consistently shown to correlate with memory parameters in all human studies.^[3]^ For instance, some studies observed that higher levels of circulating EST correlate with better working memory performance,^[25]^ whereas other studies reported changes in neither working memory nor delayed recall across the menstrual cycle.^[26]^ The mnemonic effects of EST might be task-dependent and vary with emotional valence and dose.^[27,28]^ Despite the established sex differences in memory, there is little work directly comparing EST effects in women and men.^[3]^

Sex-specific behavioural and limbic effects have also been observed for the hypothalamic peptide oxytocin.^[29,30]^ (OXT) Considering the peptide’s prosocial and anxiolytic effects, together with its high tolerability, OXT has evolved as a potential candidate compound for treating various mental disorders.^[31,32]^ Importantly, possible oxytocin-estradiol interactions have been proposed for various domains including migraine attacks^[33]^ and social anxiety.^[34]^ An intriguing notion is that both hormones may antagonistically interact in a way that yields opposing effects on limbic reactivity in women and men.^[35]^ In rodents, OXT plays a critical role in mediating social memory, with OXT receptors in the hippocampus,^[36]^ and amygdala^[37]^ being necessary for social recognition. Findings regarding OXT effects on human memory encoding have been inconsistent, with reports ranging from memory impairment^[38,39]^ to enhancement^[40–42]^ via modulation of insula activity,^[43]^ depending on inter-individual or contextual factors.^[38,44]^ Evidence for interactions between OXT and EST derives from animal studies, showing extensive coexpression of EST receptor ß in OXT neurons of the paraventricular nucleus of the hypothalamus^[45]^ and combinatorial modulation of synaptic plasticity in the medial nucleus of the amygdala in male rats.^[46]^ The EST receptor additionally binds in a dimerized form to the composite hormone response element of the OXT promotor gene and may thus induce the production of OXT.^[47–49]^ In fact, orally administered EST induced an increase in OXT plasma levels in bulimic and healthy women.^[50]^

To date, no study has simultaneously probed the modulatory mnemonic effects of both hormones and possible interactions in women and men. Therefore, we conducted a randomized, placebo-controlled, parallel-group functional magnetic resonance imaging (fMRI) study to test the effects of EST, OXT, and their interaction on emotional memory and to elucidate the neural mechanisms involved in mnestic sex differences (**see Figure 1**). Healthy men and free-cycling women were scanned under four experimental conditions: 1. transdermal placebo gel and intranasal placebo (PLC_tra_ & PLC_int_), 2. transdermal placebo and intranasal OXT (24 IU) (PLC_tra_ & OXT_int_), 3. transdermal EST_tra_ (2 mg) and intranasal placebo (EST_tra_ & PLC_int_), and 4. transdermal EST and intranasal OXT (EST_tra_ & OXT_int_). During fMRI, participants viewed positive, neutral and negative scenes. A surprise recognition task three days later was used to classify encoding trials as remembered or forgotten. We hypothesized that EST_tra_ would increase recognition memory of emotional material and activity in the hippocampus and amygdala, in both sexes. We further expected that OXT_int_ would have opposing effects in women and men (i.e., improved memory and increased insula responses in men compared to impairments and decreased insula activation in women) and that the sex-specific effects of OXT_int_ would be reduced or even inverted in the combined treatment groups.

**Figure 1.**
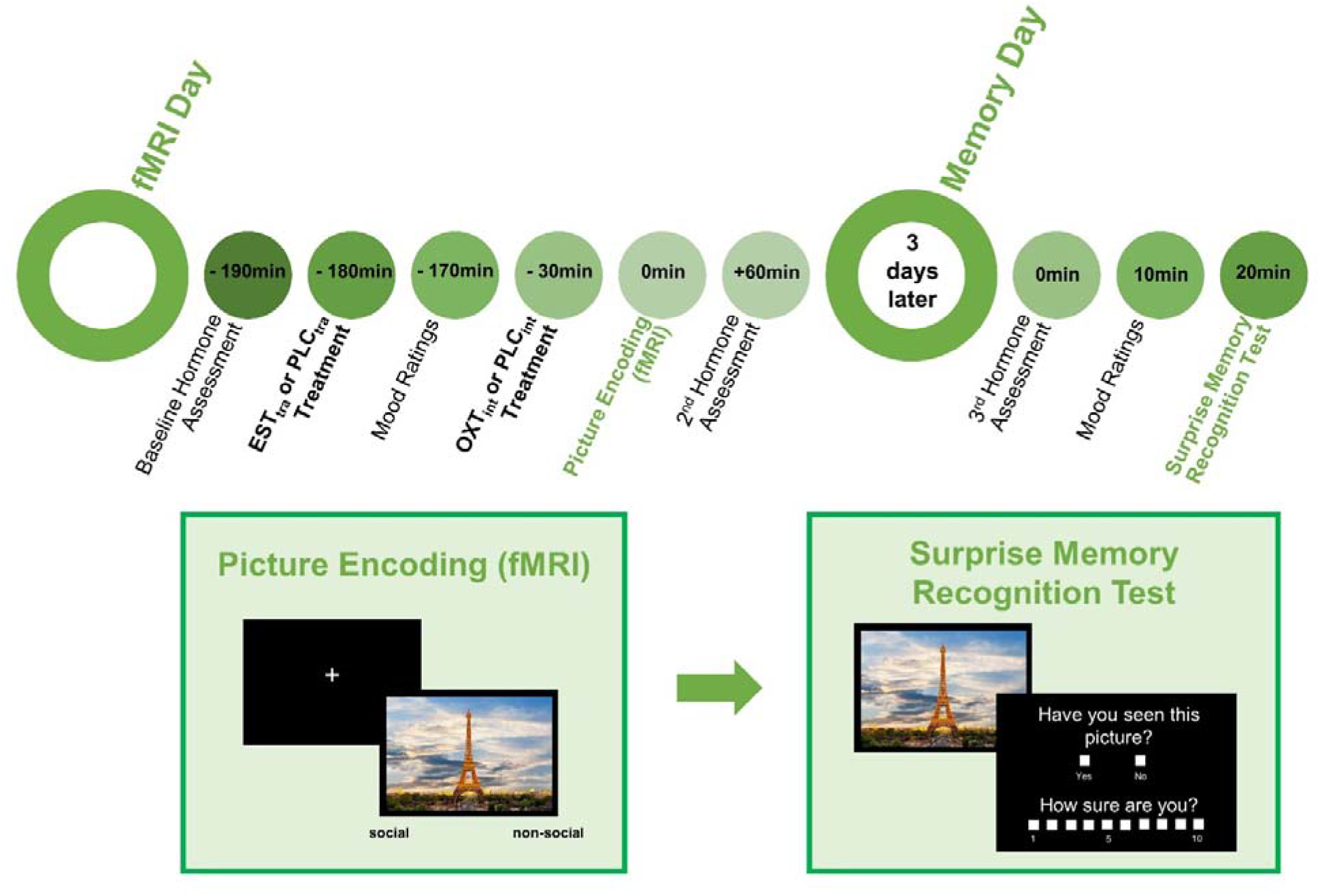
Study Design. The functional magnetic resonance imaging (fMRI) day commenced with gel administration. Nasal sprays were administered 3 hours after the gel and a high-resolution structural MRI scan was conducted 30 min after the nasal spray administration. Functional imaging data collection commenced 45 min after nasal spray treatment and included a picture encoding task. Participants viewed emotional and nonemotional scenes in a randomized order and had to press a button to indicate whether the picture’s context was social (i.e., if a human person was shown). Blood samples were collected at baseline and immediately after the fMRI testing session (approx. 4.5 hours after the gel administration). Three days later a surprise memory recognition task was administered in a second testing session and a third blood sample was collected. The recognition task included the 120 pictures shown in the scanner and 60 new distractor pictures. Participants had to rate whether they had seen the picture in the MRI task (yes/no) and their level of confidence on a 10-point Likert scale.

## Methods

### Ethics and enrolment

The study was part of a larger project and was approved by the institutional review board of the Medical Faculty of the University of Bonn and carried out in accordance with the latest revision of the Declaration of Helsinki. The study was registered in the ClinicalTrials.gov database (Identifier: NCT04330677) and the data analyses were preregistered (https://osf.io/hvknp/). The subjects were enrolled in the study after giving written informed consent, and they received a monetary reimbursement after completing the study.

### Subjects

In total, 295 subjects (160 women) were invited to a screening session prior to the testing session. The 246 subjects (122 women) who met the inclusion criteria (see below) were tested. The participants were randomly assigned to one of four experimental conditions: (1. PLC_tra_ & PLC_int_; 2. PLC_tra_ & OXT_int_; 3. EST_tra_ & PLC_int_; 4. EST_tra_ & OXT_int_). We had to exclude 20 participants from all analyses as the data of 11 subjects were not completely recorded due to technical errors and three participants did not finish the study. Additional six subjects were excluded due to hormonal (n = 4) or anatomical (n = 2) abnormalities, resulting in a sample of 226 participants (PLC_tra_ & PLC_int_: 25 men, 26 women; PLC_tra_ & OXT_int_: 33 men, 31 women; EST_tra_ & PLC_int_: 32 men, 27 women; EST_tra_ & OXT_int_: 25 men, 27 women). In accordance with our preregistered analysis of the data, we had to exclude additional 24 subjects, who remembered all or none of the stimuli in at least one valence category. Thus, our final sample for the neural and behavioural analyses included 202 subjects (PLC_tra_ & PLC_int:_ 19 men, 25 women; PLC_tra_ & OXT_int_: 27 men, 29 women; EST_tra_ & PLC_int_: 29 men, 25 women; EST_tra_ & OXT_int_: 24 men, 24 women). For demographic and psychometric baseline characteristics see **Supplementary Table S7**.

### Screening session and exclusion criteria

Screenings of the participants were conducted prior to the test sessions. Subjects were free of current or past physical or psychiatric illnesses assessed by self-report and the Mini-International Neuropsychiatric Interview.^[77]^ In addition, subjects were naive to prescription-strength psychoactive medication, and had not taken any over-the-counter psychoactive medications in the 4 weeks prior to the study. The participants were right-handed, nonsmoking and between 18 and 40 years old. Furthermore, the participants were asked to maintain their regular bed and waking times and to abstain from alcohol intake on the day of the experiment. Additional exclusion criteria were current pregnancy, MRI contraindications and the use of hormonal contraceptives. All women were tested in their early follicular phase of their menstrual cycle (days 1-6) as validated by blood assays obtained on the testing day. Female participants (n = 4) showing estradiol pre-treatment values larger than 300 pg/ml were excluded, because it can be assumed that they were not in the follicular phase of their menstrual cycle.^[78]^

### Experimental design

We used a randomized, double-blind, placebo-controlled parallel-group study design (see **Figure 1**). After a screening session, the subjects completed the fMRI testing session. Three days later a surprise memory recognition task was administered in a second testing session. The fMRI day commenced with the gel administration. In accordance with our pharmacokinetic pre-study (see **Supplementary Information** [SI]), the OXT_int_ or placebo spray was administered 3 hours after gel administration. Functional imaging data collection commenced 45 min after nasal spray administration, because it was found to be the most effective dose-test interval for OXT_int_.^[65]^ The imaging data collection included a high-resolution structural MRI scan and an emotional subsequent memory task. To validate the cycle phase and control for baseline differences in gonadal hormone levels, blood samples were collected at baseline, immediately after the fMRI testing session (approx. 4.5 hours after gel administration), and three days following the treatment.

### Treatments

#### Estradiol / placebo gel treatment

On the fMRI testing day, the subjects received either EST_tra_ gel (Estramon, 2 mg EST, Hexal AG, Holzkirchen, Germany) or placebo gel (2 mg ultrasonic gel), which was transdermally applied to the participants’ backs. In line with a pharmacokinetic study,^[79]^ a 2mg dose was chosen to reduce the possibility of side effects. The same dose has also been found to increase emotional vicarious reactivity in men when watching a distressed other.^[80]^

#### Intranasal oxytocin / placebo treatment

Via a nasal spray, the subjects self-administered 24 International Units (IU) of synthetic OXT_int_ (Sigma-Tau Industrie Farmaceutiche Riunite S.p.A., Rome, Italy) or placebo prior to the fMRI scanning in line with the standardization guidelines^[81]^ and under supervision of a trained research assistant. There is compelling evidence that OXT_int_ bypasses the blood-brain barrier and elevates OXT concentrations in the cerebrospinal fluid^[82,83]^ and brain.^[84]^ The placebo was equivalent to the OXT_int_ solution without the peptide itself. The amount of administered substance was weighed and supplemented until 24 IU were reached. The puffs were balanced between the nostrils to allow the solution to be absorbed by the nasal epithelium and an interpuff interval of approx. 45 seconds was chosen.

### Emotional subsequent memory task

#### fMRI task

During the fMRI, participants viewed a picture set (see SI) of negative (n = 40), neutral (n = 40), and positive (n = 40) scenes in a randomized order. The content of the pictures was either social (defined as the presence of a depicted human) or nonsocial, which were equally distributed across valence categories. As an attention control, the participants had to press a button, if the picture’s context was social (i.e., if a human person was shown). The participants could choose their responses using an MRI-compatible response grip system (NordicNeuroLab AS, Bergen, Norway). The paradigm was written in Presentation code (Neurobehavioral Systems, Albany, CA) and the stimuli were presented on a 32-inch MRI-compatible TFT LCD monitor (NordicNeuroLab, Bergen, Norway) placed at the rear of the magnet bore.

#### Surprise recognition task

The participants performed a surprise memory recognition task three days after the MRI scan to classify pictures as remembered and nonremembered. The recognition task included the 120 pictures shown in the scanner and 60 new distractor pictures (matched for valence and arousal ratings, see **SI**). The participants had to rate whether they had seen the picture in the MRI task (yes/no) and their level of confidence on a 10-point Likert scale.

### Data analysis

#### fMRI data acquisition and analysis

All fMRI data were acquired using a 3T Siemens TRIO MRI system (Siemens AG, Erlangen, Germany) with a Siemens 32-channel head coil. fMRI data were preprocessed and analysed using standard procedures in SPM12 software (Wellcome Trust Center for Neuroimaging; http://www.fil.ion.ucl.ac.uk/spm) implemented in MATLAB (MathWorks). Six conditions (valence (3) * memory (2)) were modelled by a stick function convolved with a hemodynamic response function. Button presses were included as regressors of no interest. On the first level, task-specific effects were modelled. On the second level, a full factorial design with the between-subjects variables “OXT_int_ treatment”, “EST_tra_ treatment”, and “sex” was conducted. Based on previous findings ^[10,11,16,43]^, analyses were conducted focusing on the anatomically defined amygdala, hippocampus and insula cortex based on the WFU PickAtlas as regions of interest (ROIs). The significance threshold for the ROI analyses was set to *p* < 0.05, familywise error-corrected for multiple comparisons based on the size of the ROI. In addition, an exploratory whole-brain analysis was performed (cluster defining threshold *p* < 0.001; significance threshold *p*_FWE_ < 0.05 at peak level). Parameter estimates of significant contrasts were extracted using MarsBaR (https://www.nitrc.org/projects/marsbar, RRID: SCR_009605) and further analszed in SPSS 25 (IBM Corp., Armonk, NY). For further analyses and details see SI.

#### Memory performance

Stimuli were classified as remembered if the picture was included in the emotional subsequent memory task and correctly identified in the recognition task. In accordance with our preregistration, the participants had to rate their confidence of the classification for an item as ≥ 2 on the 10-point Likert scale to be classified as remembered. The participants who remembered all or none of the stimuli in at least one valence category were excluded from the analysis (n = 24), because a minimum of one trial was required in every valence category for the neural and behavioral model estimation. We calculated d prime (*d‘*) by subtracting the z-standardized false alarm rate from the z-standardized hit rate. The hit rate was the mean of the correctly identified stimuli used in the fMRI paradigm. The false alarm rate was the mean of the distractors in our recognition paradigm, which were incorrectly identified as seen by the subject, although they were not included in the fMRI task. When the hit rate or false alarm rate equals zero or one, the corresponding z-score would be −/+∞. Thus, we adjusted *d‘* according to the loglinear method ^[85]^ by adding 0.5 to both the number of hits and the number of false alarms and adding 1 to both the number of signal trials and noise trials, before calculating the hit and false alarm rates.^[86]^ A high *d‘* indicated that the signal was easily detected.^[87]^

### Statistical analyses

Behavioural, neuroendocrine, and demographic data were analysed in SPSS 25 using standard procedures including analyses of variances (ANOVAs) and post-hoc *t*-tests. Post hoc *t*-tests were Bonferroni-corrected (*p*_cor_).In general, two-tailed *p* values < .05 were considered significant, except for directional hypotheses, in which case one-tailed t-tests were calculated. If the assumption of sphericity was significantly violated, a Greenhouse-Geisser correction was applied. As measures of effect sizes, partial eta-squared and Cohen’s d were calculated. Furthermore, frequentist inference was complemented by computing Bayes factors (BFs) with default priors via JASP (version 0.14.1.0). For Bayesian *t*-tests, BF_10_ is reported. A BF_10_ larger than 1 can be interpreted as evidence in favour of the alternative hypothesis given the current data. For Bayesian ANOVAs, BF_incl_ (with fixed seed = 1) was calculated, which compares the performance of all models that include the effect to the performance of all the models that do not include the effect. Mixed-design ANOVAs with between-subjects variables “OXT_int_ treatment” (OXT_int_, placebo nasal spray), “EST_tra_ treatment” (EST_tra_, placebo gel) and “sex” (women, men) and the within-subject factors “valence” (negative, neutral, positive) and “sociality” (social, nonsocial) were conducted for the outcome “memory” (*d’*). In a second and third model, either the factor “sociality” or the factor “valence” was aggregated. In a fourth model, both factors “valence” and “sociality” were aggregated to provide an overall *d’* that represented the general memory performance of the participants irrespective of valence and sociality. Changes in hormone concentrations were examined with mixed-design ANOVAs with the between-subject factors “OXT_int_ treatment”, “EST_tra_ treatment”, and “sex” and the within-subject variable “time” (baseline, after treatment, and three days after treatment; for OXT changes: baseline vs. after treatment). Furthermore, to explore the potential moderating effects of treatment-induced hormonal changes, the magnitude of the increases in hormone concentrations (levels of EST, OXT, testosterone, and progesterone after the fMRI session minus baseline) were considered covariates in the main analyses with significant behavioural (*d’*) and neural outcomes (i.e., parameter estimates of significant contrasts of interests).

## Results

### Neuroendocrine parameters

At baseline, women had significantly higher EST concentrations (*t*_(117.24)_ = −5.70, *p* < 0.001, *d* = −0.80; BF_10_ = 206050.22), but lower testosterone (*t*_(98.31)_ = 27.09, *p* < 0.001, *d* = 3.89; BF_10_ = 9.96 x 10^64^) and OXT levels (*t*_(196)_ = 2.75, *p* < 0.01, *d* = 0.39; BF_10_ = 5.04) than men. The progesterone baseline concentrations were comparable between the two sexes (*t*_(98.72)_ = − 1.76, *p* = 0.08, *d* = −0.25; BF_10_ = 0.65). Importantly, all baseline levels were comparable between treatment groups in women and men (all *p*s > 0.05; BF_10_ < 1).

The EST_tra_ administration significantly increased blood EST levels in both sexes (see **Figure 2 and Supplementary Table S4**; time * EST_tra_ treatment: *F*_(1.00, 182.64)_ = 265.92, *p* < 0.001, η_p_^2^ = 0.60; BF_incl_ = 1.14 * 10^79^), with women exhibiting a significantly larger increase than men (time * sex * EST_tra_ treatment: *F*_(1.01, 182.645)_ = 16.30, *p* < 0.001, η_p_ ^2^ = 0.08; BF_incl_ = 742418.03). There were no significant main or interaction effects of the OXT_int_ treatment on EST levels (all *p*s > 0.05; BF_10_ < 0.2), indicating that the OXT_int_ treatment did not modulate the EST increase.

**Figure 2.**
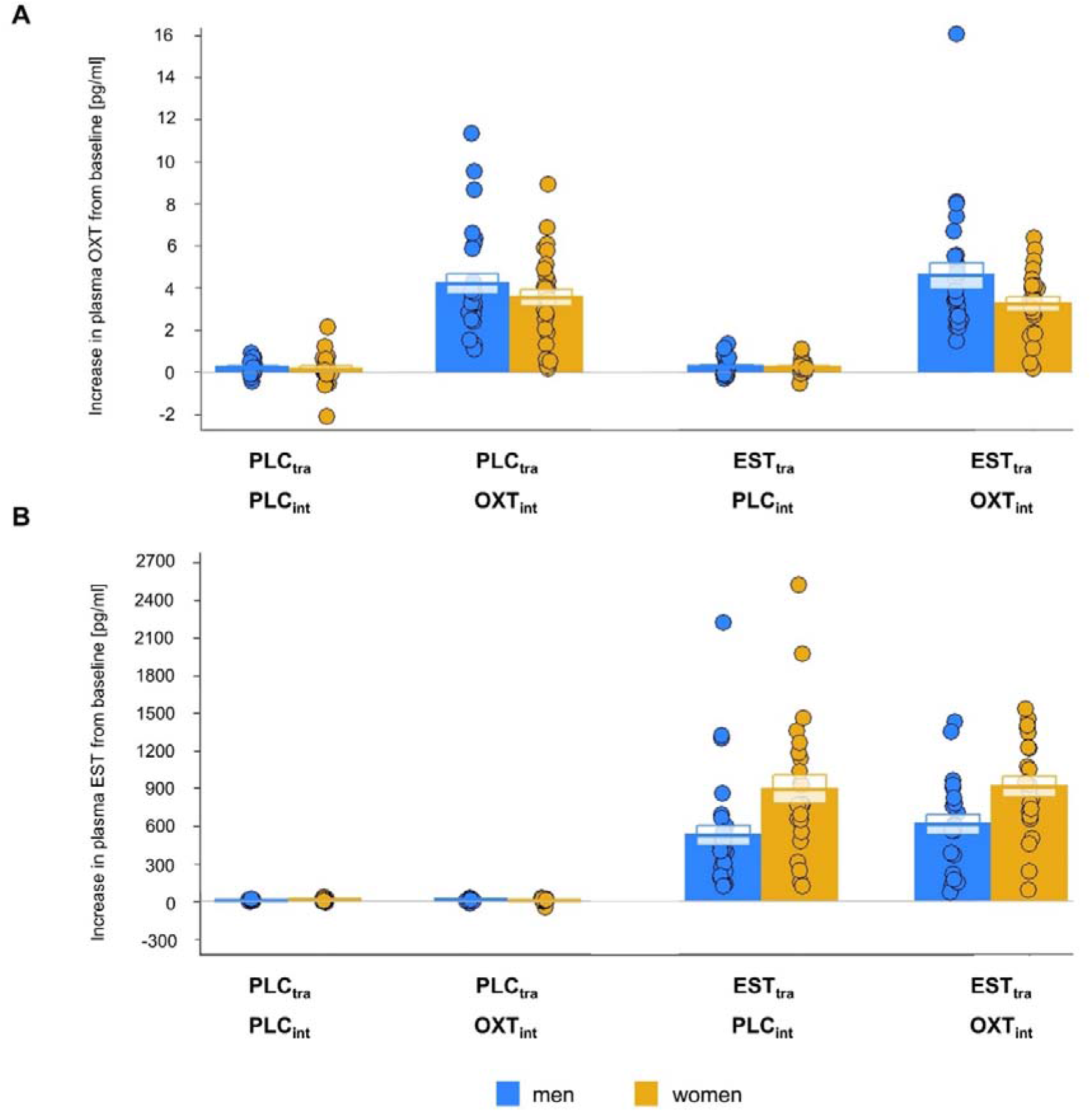
Treatment-induced changes (immediately after fMRI session minus baseline) in oxytocin and estradiol plasma levels. **A** The administration of 24 international units (IU) of intranasal (OXT_int_) induced a significant increase in blood oxytocin levels in both sexes. There was no significant interaction with the estradiol treatment. **B**The transdermal administration of 2 mg estradiol (EST_tra_) significantly elevated blood estradiol levels in both sexes. EST_tra_ treatment induced a significantly stronger increase in women than in men, but there was no significant interaction with oxytocin treatment. PLC_tra_ = transdermal placebo gel; PLC_int_ = intranasal placebo; OXT_int_ = intranasal oxytocin; EST_tra_ = transdermal estradiol.

As an additional control analysis, we used a median-dichotomization and excluded EST_tra_-treated women with large EST increase. In this subsample, the treatment-induced increases in EST levels were comparable between women and men within the treatment groups (all *p*s > 0.05; BF_10_ < 0.6) and the main behavioral and neural analyses yielded a similar pattern of results (see **SI** and **Supplementary Figure S1**).

OXT_int_ administration significantly increased blood oxytocin levels in both sexes (see **Figure 2 and Supplementary Table S5**; time * OXT_int_ treatment: *F*_(1,190)_ = 215.77, *p* < 0.001, η_p_^2^ = 0.53; BF_incl_ = 2.43 * 10^31^) with men exhibiting a nonsignificantly larger increase than women (time * sex * OXT_int_ treatment: *F*_(1, 190)_ = 3.42, *p* = 0.07, η_p_^2^ = 0.02; BF_incl_ = 1.00). There were no significant main or interaction effects of the EST_tra_ treatment on the OXT levels (all *p*s > 0.05; BF_10_ < 0.4), indicating that the EST_tra_ treatment did not modulate the OXT increase.

To examine whether the changes in OXT_int_ and EST_tra_ levels moderated behavioural and neural treatment effects, we included the hormonal changes (after treatment minus baseline) as separate covariates in the analyses and all sex * treatment interactions remained significant. Furthermore, the treatments did not significantly alter the participants’ mood (see **SI** and **Supplementary Table S6**).

### Behavioural results

Recognition memory was significantly better with emotional than neutral items (main effect valence: *F*_(2,388)_ = 13.18, *p* < 0.001, η_p_^2^ = 0.06; BF_incl_ = 3054.02) and with social than nonsocial stimuli (main effect of sociality: *F*_(1,194)_ = 7.59, *p* < 0.01, η_p_^2^ = 0.04; BF_incl_ = 5.29). However, there were no significant interactions between these factors (valence and sociality) and sex or treatments (BF_incl_ < 0.4). Thus, the following analyses are based on *d’* averaged across valences and sociality.

#### Behavioural results: sex differences

Sex differences in recognition memory were significantly altered by both EST_tra_ and OXT_int_ treatments (significant three-way interaction: sex * EST_tra_ treatment * OXT_int_ treatment: *F*_(1,194)_ = 6.96, *p* < 0.01, η_p_ ^*2*^ = 0.04; BFincl = 5.55; see **Figure 3 and Supplementary Table S1**). Under placebo (PLC_tra_ & PLC_int_), women showed significantly better memory performance than men (*t*_(42)_ = −2.96, *p* < 0.01 [*p*_cor_ = 0.02], *d* = −0.90; BF_10_ = 8.31), but there were no significant sex differences in the EST_tra_ & PLC_int_ group (*t*_(52)_ = −0.32, *p* = 0.75, *d* = −0.09; BF_10_ = 0.29) or the PLC_tra_ & OXT_int_ group (*t*_(54)_ = 0.21, *p* = 0.83, *d* = 0.06; BF_10_ = 0.28). After the combined treatment (EST_tra_ & OXT_int_), a trend similar to that observed in the placebo group was evident (women > men; *t*_(46)_ = −1.87, *p* = 0.07 [*p*_cor_ = 0.28], *d* = −0.54; BF_10_ = 1.16). Thus, the Bayes factors indicated moderate evidence for sex differences under placebo, moderate evidence for the absence of sex differences after single treatments and an inconclusive sex effect in the combined group.

**Figure 3.**
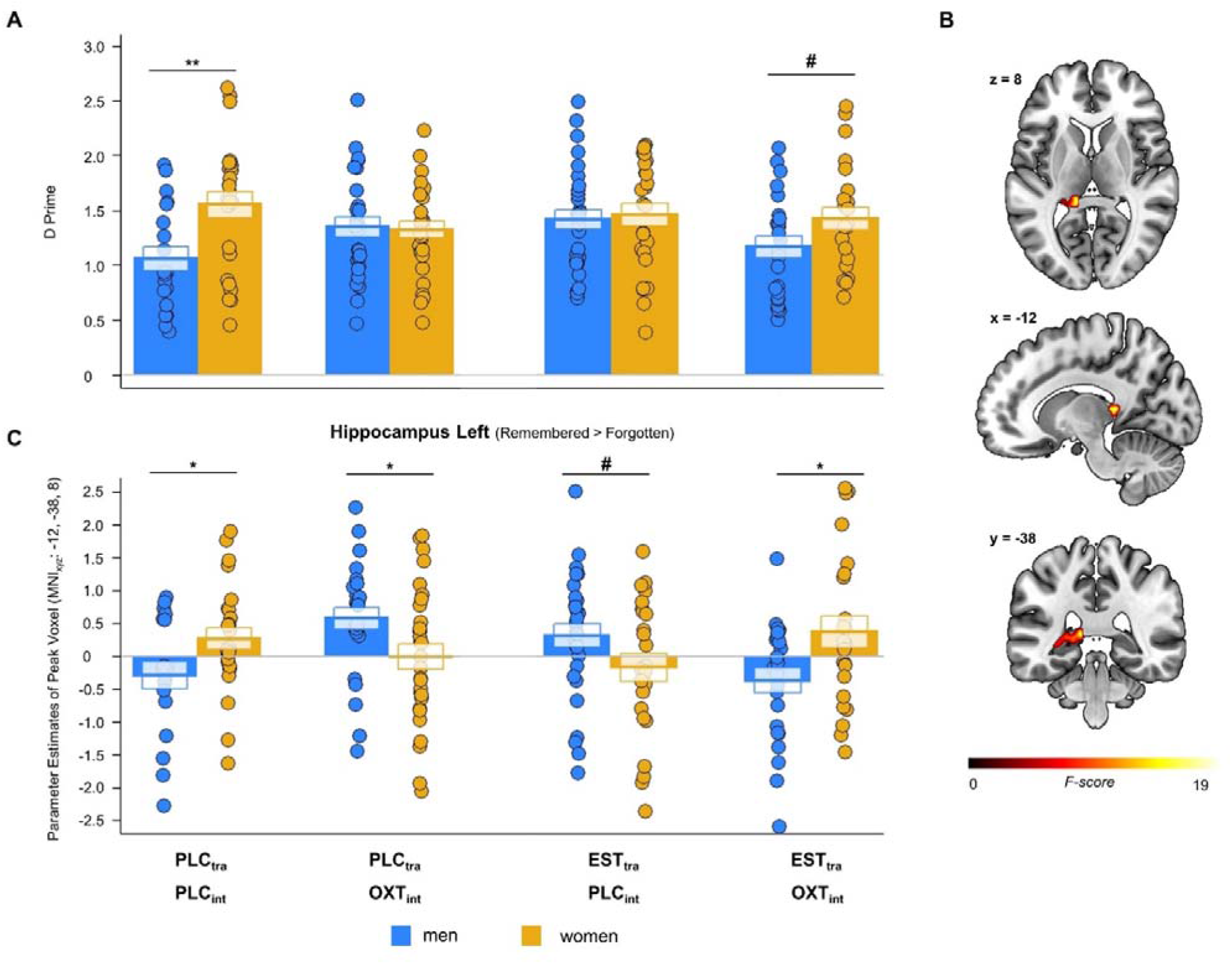
Treatment effects on mnemonic and hippocampal sex differences. **A** Following placebo administration, women showed a significantly better memory performance (*d’*) than men, but there were no significant sex differences after single trandermal estradiol (EST_tra_) or intranasal oxytocin (OXT_int_) treatment. After the combined treatment, a trend similar to that observed in the placebo group was evident. **B**We observed a significant three-way interaction of sex, OXT_int_, and EST_tra_ treatment in the left hippocampus responses to remembered compared to forgotten stimuli (MNI peak coordinates [x, y, z]: −12, −38, 8). **C** Further analyses of the parameter estimates for the left hippocampal responses revealed a similar pattern as that observed with memory performance. Women exhibited stronger left hippocampal responses to remembered items compared to forgotten items than men under placebo. This sex difference was reversed after either EST_tra_ or OXT treatment. Intriguingly, the same pattern as that observed in the placebo group was again evident in the combined treatment group. PLC_tra_ = transdermal placebo gel; PLC_int_ = intranasal placebo; OXT_int_ = intranasal oxytocin; EST_tra_ = transdermal estradiol. ^#^*p* < 0.1, \**p* < 0.05, \*\**p* < 0.01.

#### Behavioural results: treatment effects

To disentangle the observed three-way interaction, we examined the treatment effects separately for both sexes. We found a significant interaction between the OXT_int_ and EST_tra_ treatments in men (*F*_(1,95)_ = 7.84, *p* < 0.01 [*p*_cor_ = 0.01], η_p_^*2*^ = 0.08; BF_incl_ = 6.94), but not in women (*F*_(1,99)_ = 0.93, *p* = 0.34, η_p_ ^*2*^ = 0.01; BF = 0.40). In men, OXT_int_ nonsignificantly improved recognition memory after PLC_tra_ treatment (PLC_tra_ & OXT_int_ vs. PLC_tra_ & PLC_int_: *t*_(44)_ = −2.01, *p* = 0.03 [*p_cor_* = 0.1], *d* = −0.6; BF_10_ = 1.45; one-tailed) and reduced performance after EST_tra_ treatment (EST_tra_ & OXT_int_ vs. EST_tra_ & PLC_int_: *t*_(51)_ = 1.95, *p* = 0.06 [*p*_cor_ = 0.23], *d* = 0.54; BF_10_ = 1.30). Likewise, in men, EST_tra_ treatment produced significantly better memory in participants who received PLC_int_ (EST_tra_ & PLC_int_ vs. PLC_tra_ & PLC_int_: *t*_(46)_ = −2.55, *p* < 0.01 [*p*_*cor*_ = 0.03], *d* = −0.75; BF_10_ = 3.75; one-tailed) and an impairment in individuals who received OXT_int_ (EST_tra_ & OXT_int_ vs. PLC_tra_ & OXT_int_: *t*_(49)_ = 1.37, *p* = 0.18, *d* = 0.38; BF_10_ = 0.60). Together, the Bayes factors indicate moderate evidence for an EST_tra_ * OXT_int_ interaction in men and moderate evidence for an EST_tra_-induced memory improvement in men who had received PLC_int_.

### Neural results: task effects

Across treatments and sexes, the remembered items compared to forgotten items induced activations in a wide network of brain areas (see **Supplementary Table S2**), including the bilateral hippocampus (left: Montreal Neurologocal Institute [MNI] peak coordinates [x, y, z]: −32, −18, −14, *F*_(1,194)_ = 28.81, on peak level *p*_FWE_ < 0.02; right: MNI peak coordinates [x, y, z]: 20, −6, −14, *F*_(1,194)_ = 105.12, on peak level *p*_FWE_ < 0.001). Furthermore, an emotional memory effect (i.e., [Emotional _Remembered > Forgotten_ > Neutral _Remembered > Forgotten_]) was evident in the left and right amygdala (left: MNI peak coordinates [x, y, z]: −20, −6, −14, *F*_(1,194)_ = 18.12, on peak level *p*_FWE_ < 0.01; right: MNI peak coordinates [x, y, z]: 22, −6, −12, *F*_(1,194)_ = 13.96, on peak level *p*_FWE_ = 0.02), as well as the left hippocampus (MNI peak coordinates [x, y, z]: −18, −6, −14, *F*_(1,194)_ = 18.55, on peak level *p*_FWE_ < 0.01) and the right insula (MNI peak coordinates [x, y, z]: 26, 22, −16, *F*_(1,194)_ = 18.85, on peak level *p*_FWE_ = 0.01; for additional activations, see **Supplementary Table S3**).

### Neural results: sex differences

We observed a significant main effect of sex in the left amygdala responses to positive remembered stimuli compared to positive forgotten stimuli (MNI peak coordinates [x, y, z]: −28, −2, −24, *F*_(1,204)_ = 12.37, on peak level *p*_FWE_ = 0.03; see **Figure 4**) and in the right amygdala responses to negative remembered stimuli compared to negative forgotten stimuli (MNI peak coordinates [x, y, z]: 28, 4, −16, *F*_(1,205)_ = 12.23, on peak level *p*_FWE_ = 0.03), but there were no significant interactions between sex and either treatment type. Analyses of the extracted parameter estimates revealed that across treatments men exhibited significantly increased right amygdala responses to negative remembered stimuli relative to negative forgotten stimuli compared to women (*t*_(200)_ = 3.16, *p* < 0.01, *d* = 0.45; BF_10_ = 15.17). In contrast, women showed a stronger activation than men in response to positive remembered stimuli relative to positive forgotten stimuli in the left amygdala (*t*_(200)_ = −2.85, *p* < 0.01, *d* = − 0.40; BF_10_ = 6.50).

**Figure 4.**
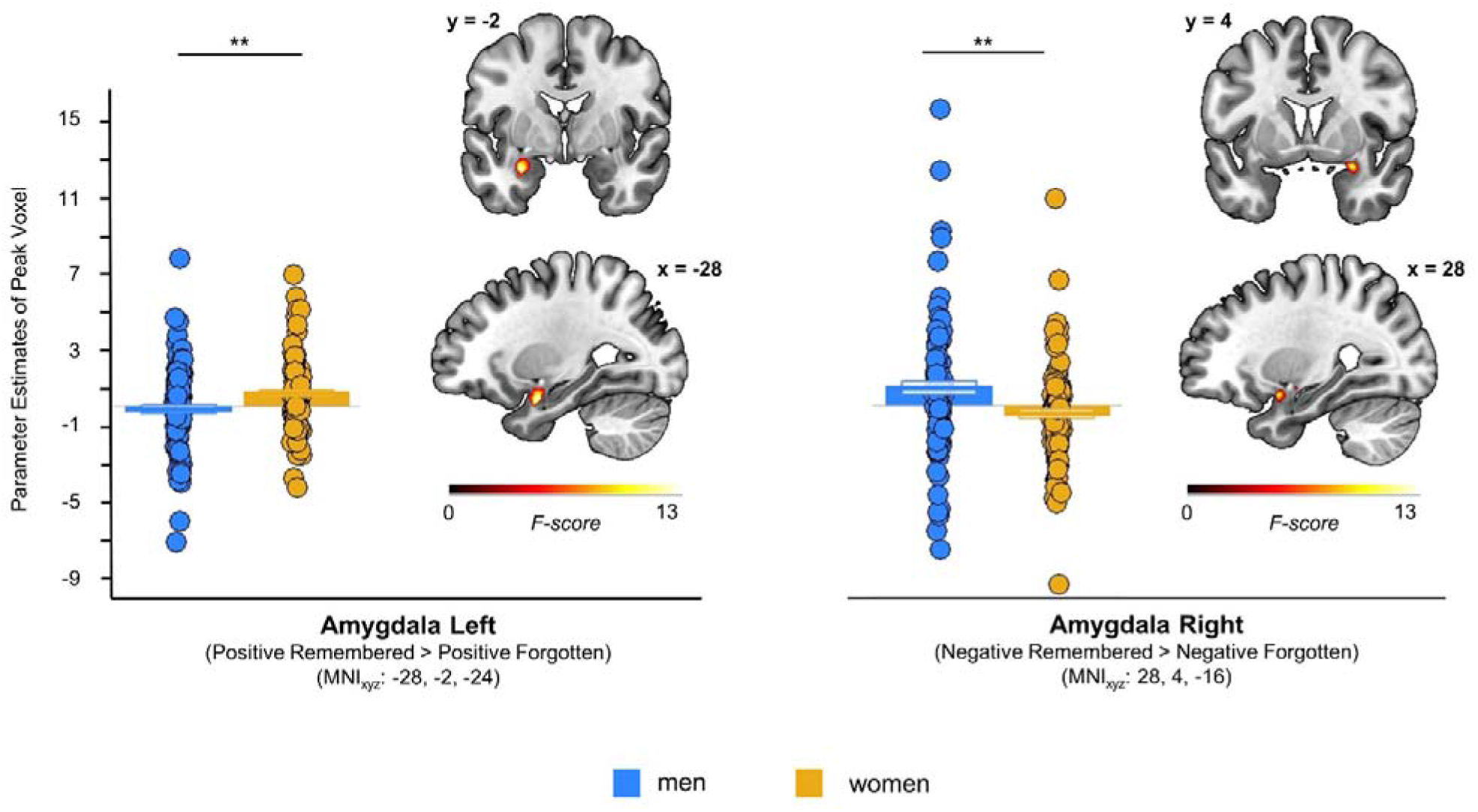
Sex-specific lateralization of amygdala activation. Significant main effects of sex emerged in left amygdala responses to positive remembered stimuli compared to positive forgotten stimuli (MNI peak coordinates [x, y, z]: −28, −2, −24) and for right amygdala responses to negative remembered stimuli compared to negative forgotten stimuli (MNI peak coordinates [x, y, z: 28], 4, −16). Across treatments, analyses of the extracted parameter estimates showed that men exhibited significantly greater right amygdala responses to negative remembered stimuli relative to negative forgotten stimuli than women. However, the pattern in the left amygdala was reversed. Women showed a stronger activation than men in response to positive remembered stimuli compared to positive forgotten stimuli. There were no significant interactions between sex and either treatment type. \*\**p* < 0.01.

Furthermore, we found a significant three-way interaction of sex, OXT_int_, and EST_tra_ treatment in the left hippocampus responses to remembered stimuli compared to forgotten stimuli (MNI peak coordinates [x, y, z]: −12, −38, 8, *F*_(1,194)_ = 17.80, on peak level *p*_FWE_ = 0.012; see **Figure 3**). Further analyses of the extracted parameter estimates revealed that women exhibited nonsignificantly stronger hippocampal responses to remembered items than men under placebo (PLC_tra_ & PLC_int_; *t*_(42)_ = −2.10, *p* = 0.04 [*p_cor_* = 0.17], *d* = −0.64; BF_10_ = 1.68). This sex difference was reversed (i.e., men > women) in both the EST_tra_ & PLC_int_ group (*t*_(52)_ = 1.77, *p* = 0.08, [*p_cor_* = 0.33], *d* = 0.48; BF_10_ = 0.99) and the PLC_tra_ & OXT_int_ group (*t*_(54)_ = 2.25, *p* = 0.03 [*p_cor_* = 0.11], *d* = 0.60; BF_10_ = 2.13). Interestingly, the same pattern observed in the placebo group was again evident in the combined treatment group (women > men; *t*_(46)_ = −2.42, *p* = 0.02 [*p_cor_* = 0.08], *d* = −0.7; BF_10_ = 2.93). Notably, a similar sex * EST_tra_ treatment * OXT_int_ treatment interaction emerged for the emotional memory effect in the left hippocampus ([Emotional _Remembered > Forgotten_ > Neutral _Remembered > Forgotten_]; MNI peak coordinates [x, y, z]: −14, −40, 10, *F*_(1,194)_ = 16.66, on peak level *p*_FWE_ = 0.02; see Supplement). We did not observe a significant three-way interaction of sex, OXT_int_, and EST_tra_ treatment in the insula.

### Neural results: treatment effects

To disentangle the observed three-way interaction in the left hippocampus, we examined the parameter estimates for left hippocampal responses to remembered items compared to forgotten items separately for the both sexes. We found a significant interaction between the OXT_int_ and EST_tra_ treatments in men (*F*_(1,95)_ = 17.30, *p* < 0.001 [*p*_cor_ < 0.001], η_p_^2^ = 0.15; BF_incl_ = 316.43), but not in women (*F*_(1,99)_ = 3.92, *p* = 0.05 [*p*_cor_ = 0.10], η_p_^2^ = 0.04; BF_incl_ = 1.38). OXT_int_ significantly increased the hippocampal response to remembered stimuli compared to forgotten stimuli in men who had received PLC_tra_ (PLC_tra_ & OXT_int_ vs. PLC_tra_ & PLC_int_: *t*_(44)_ = − 3.30, *p* < 0.01 [*p*_cor_ < 0.01], *d* = −0.99; BF_10_ = 18.29), but had the opposite effect in men after EST_tra_ treatment (EST_tra_ & OXT_int_ vs. EST_tra_ & PLC_int_: *t*_(51)_ = 2.61, *p* = 0.01 [*p*_cor_ = 0.04], *d* = 0.72; BF_10_ = 4.24). Likewise, EST_tra_ nonsignificantly increased hippocampal activation in men after PLC_int_ (EST_tra_ & PLC_int_ vs. PLC_tra_ & PLC_int_: *t*_(46)_ = −2.17, *p* = 0.02 [*p*_cor_ = 0.07], *d* = −0.64; BF_10_ = 1.88; one-tailed) and produced significant effects in the opposite direction when combined with OXT_int_ treatment (EST_tra_ & OXT_int_ vs. PLC_tra_ & OXT_int_: *t*_(49)_ = 3.80, *p* < 0.001 [*p*_cor_ < 0.001], *d* = 1.07; BF_10_ = 67.05). Thus, again, the Bayes factors indicated strong evidence for an EST_tra_ * OXT_int_ interaction in men. Furthermore, there was moderate-to-strong evidence for opposing effects of OXT_int_ on hippocampal activation depending on EST_tra_ pretreatment and strong evidence for an EST_tra_-induced decrease when combined with OXT_int_.

## Discussion

The goal of the current study was to elucidate the effects of EST_tra_ and OXT_int_ and their interaction on episodic memory in healthy women and men. Our results revealed that women showed better recognition memory than men and increased hippocampal responses to subsequently remembered items irrespective of emotional valence. Separate treatments with either EST_tra_ or OXT_int_ significantly diminished this mnemonic sex difference and reversed the hippocampal activation pattern. However, the combined treatments led to a memory performance comparable to that of the placebo group, indicating an antagonistic effect of the two hormones at the administered doses. This pattern was also evident in a subsample with comparable treatment-induced EST increases in women and men. Given significant memory differences between the sexes in the placebo group, our data are consistent with the reported advantage of women in episodic memory, ^[1,3,5–8]^ which may contribute to the higher prevalence of stress- and trauma-related disorders in women. The preponderance of studies with exclusively female samples to examine the effects of the “female” sex hormone EST_tra_ and exclusively male samples in the case of OXT_int_ might have obfuscated the contribution of these hormones to sex differences in episodic memory.^[16,51]^ Our study included both sexes and indicated an essential role of EST_tra_ and OXT_int_ as modulators of episodic memory in women and men.

In men, separate treatment with either EST_tra_ or OXT_int_ increased hippocampal activation and improved episodic memory. These findings are in line with previous studies showing that EST and OXT receptor signalling in the hippocampus regulate neuronal excitability, synaptic plasticity, and memory formation in male mice and rats.^[52,53]^ For instance, EST induced spinogenesis in the hippocampus^[54]^ and improved long-term memory following pretraining administration of EST_tra_ in male rats.^[55,56]^ Likewise, OXT has been found to enhance cortical information transfer in the hippocampus irrespective of sex by exciting fast-spiking interneurons.^[57]^ Furthermore, conditional deletion of OXT receptors in the hippocampus of male mice impaired the persistence of long-term social recognition memory,^[58]^ and OXT_int_ rescued recognition memory and hippocampal long-term potentiation in stressed male rats.^[59]^ We are not aware of any study probing the effects of exogenous EST_tra_ on memory performance in men, but several studies found improved recognition memory after OXT_int_ administration^[40,41,60]^ (but also see ^[38,61]^). We did not observe a selective facilitation of social stimuli or valence-dependent effects on insula reactivity, which could be related to the use of scenes with a human person as social items instead of faces and may reflect differences between recognition and recall memory.^[43]^ Selective OXT_int_ effects in men, but not in women, have been previously observed for hippocampal responses to cooperative interactions^[29]^ and chemosensory stress signals.^[62]^ Furthermore, OXT_int_ appears to modulate hippocampus activation as a function of social salience, with the peptide decreasing hippocampal responses to an unfamiliar child, but not the own child in fathers.^[63]^ Intriguingly, a single EST_tra_ treatment produced similar effects as OXT, but the OXT_int_-induced increase in hippocampal activation was absent after EST_tra_ pretreatment. One possible explanation for this pattern of results is that EST_tra_ may have increased OXT receptor binding,^[64]^ thereby mirroring the opposing effects previously observed for higher OXT_int_ doses in men.^[65]^ EST_tra_ treatment did not trigger the release of endogenous OXT_int_ 4.5 hours after gel administration, but there is preliminary evidence that EST_tra_-induced OXT secretion would be most pronounced 18-36 hours later.^[50]^ While we detected no significant difference between treatment groups in EST and OXT concentrations before the surprise recognition task 3 days after the fMRI session, elevated hormone levels may have influenced both encoding and early consolidation. Interestingly, EST_tra_*OXT_int_ interactions may have contributed to the modulatory effects of hormonal contraception^[66]^ and to the sex-specific effects of OXT_int_ that have been found in various domains, including fear-related amygdala reactivity^[35]^, the perception of competition,^[67]^ moral decision-making,^[30]^ and emotional responses to couple conflict.^[68]^

The absence of a significant EST_tra_ effect in women may reflect sex-specific dose-dependent mechanisms. The EST_tra_ treatment produced a significantly larger increase in peripheral EST concentrations in women than in men, yielding concentrations comparable to the EST levels in pregnancy.^[69]^ Thus, the nonsignificant decrease in hippocampal activation is consistent with the notion of an inverted U-shaped dose-response function of EST_tra_ in women.^[27]^ EST_tra_ levels within physiological ranges have been found to stimulate hippocampal activity, while levels within supraphysiological ranges can have the opposite effect. While low levels of EST were shown to activate the high-affinity receptor ERα, thus inducing synaptogenesis and enhancing blood oxygen level-dependent (BOLD) responses, high levels of EST also activated the low-affinity receptor ERβ, which in turn reduced synaptogenesis.^[70,71]^ As such, the EST_tra_-enhanced anxiolytic action of OXT_int_ previously observed in female mice may also be dose-dependent.^[48]^ In contrast, sex-specific effects of OXT_int_ on the amygdala response to fearful faces in women have been reported across a range of doses,^[35]^ but OXT_int_ effects on episodic memory have not been systematically tested in women. Intranasal OXT administered after acquisition improved recognition memory of faces in a combined sample of 36 women and men,^[42]^ suggesting that the peptide’s mnemonic effect may differ between acquisition and encoding or may be modulated by the stimulus material.

Interestingly, we found the expected memory advantage for emotional material and replicated the sex-specific lateralization of amygdala recruitment,^[9,12–14]^ but there was no significant interaction of the treatments for the emotional memory effect in the amygdala. The effects of EST_tra_ and OXT_int_ on amygdala activation have been well established in animal studies,^[46,72,73]^ but translation to humans may depend on methodological details such as task design and stimulus material. Furthermore, while the availability of aromatase, the enzyme that catalyzes testosterone to estradiol, is comparable in the amygdala between women and men, higher aromatase availability was associated with lower memory performance in men, but not women.^[74]^ Thus, the effects of and interactions with other gonadal steroids should be taken into consideration. For instance, testosterone treatment shifted amygdala lateralization towards the right hemisphere in transgender boys^[75]^ and exogenous progesterone increased amygdala responses to emotional faces in women.^[76]^

The findings of the present study need to be considered in the context of the following limitations. In both sexes, supraphysiological estradiol levels were induced due to the exogenous estradiol administration. It is conceivable that interactions between OXT_int_ and EST_tra_ in women would be evident at physiological EST levels occurring during the menstrual cycle. Future studies should employ different doses and postlearning administration protocols to further delineate the sex-specific memory effects of EST_tra_ and OXT_int_. Additional clinical trials using long-term applications are needed to further disentangle the hormones’ impact on memory formation and to deduce possible direct implications for the development of several psychiatric disorders, such as PTSD.

Collectively, our results provide evidence that EST_tra_ and OXT_int_ modulate episodic memory and hippocampal functioning in men. Concomitant pharmacological enhancement of EST_tra_ and OXT_int_ signalling produced antagonistic effects that may contribute to the previously observed sex-specific hormonal effects in various domains. Our findings support the increasingly recognized notion that it is vital to consider sex differences in pharmacological clinical trials.

## Acknowledgements

The authors thank Mitjan Morr and Jana Lieberz for helpful discussions about our manuscript.

## Study approval statement

This study protocol was reviewed and approved by the institutional review board of the medical faculty of the University of Bonn [Approval number: 213/16]. Written informed consent was obtained from all participants included in this study.

## Competing interests

The authors report no competing biomedical financial interests or personal affiliations in connection with the content of this manuscript.

## Role of funding source

R.H. and D.S. are supported by a German Research Foundation (DFG) grant (HU 1302/11-1 and SCHE 1913/5-1). The DFG had no further role in study design; in the collection, analysis and interpretation of data; in the writing of the report; and in the decision to submit the paper for publication.

## Author Contributions

M.C., and D.S. designed the experiment; M.C., I.T. and B.A. conducted the experiments; M.C., B.S-W. and D.S. analysed the data. M.C. and D.S. wrote the manuscript. All authors read and approved the manuscript in its current version.

## Data availability statement

A preprint of this article is published at the BioRxiv preprint server (https://doi.org/10.1101/2021.11.22.469500). The data that support the findings of the present study are openly available in the repository of the Open Science Foundation at https://osf.io/hvknp/ (doi: 10.17605/OSF.IO/HVKNP). The unthresholded statistical maps of the fMRI results can be accessed at https://neurovault.org/collections/FBHLSKJX/. The code that supports the findings of the present study is openly available in the repository of the Open Science Foundation at https://osf.io/hvknp/ (doi: 10.17605/OSF.IO/HVKNP).

## Notes

https://osf.io/hvknp/

https://neurovault.org/collections/FBHLSKJX/

### Competing Interest Statement

The authors have declared no competing interest.

### Summary of Updates

Abstract updated. Minor Updates in the Main Manuscript for clarification. Supplemental files updated and additional Figure added.

