## Supplementary Material for "Exogenous estradiol and oxytocin modulate sex differences in hippocampal reactivity and episodic memory"

**Supplementary Methods**

**Pharmacokinetic pre-study**

We conducted a prestudy involving 10 healthy participants (5 women; mean age ± SD = 24.10 ± 4.07 years), to examine the pharmacokinetics of estradiol gel (Estramon 2 mg estradiol, Hexal AG, Holzkirchen, Germany) administration. Blood samples were taken prior to estradiol administration (i.e., baseline) and in 1-hour intervals after drug application up to 5 hours post administration. An additional blood sample was taken the next day (after 18 hours). Serum estradiol levels peaked 3-4 hours after gel administration, but a significant increase relative to baseline was already evident after 2 hours (*t*_(9)_ = 2.44, *p* = 0.04, *d* = 1.10; **cf.**^[1]^). Estradiol levels remained significantly elevated throughout the last measurement. A previous study tested the topical administration of a different drug (Divigel, Orion Pharma AG, Zug, Switzerland) containing 2 mg estradiol and found significantly increased estradiol serum concentrations as soon as 1 hour after administration and maximum average levels after 2 hours.^[2]^

**Power analysis**

We used G*Power 3^[3]^ to conduct an a-priori power analysis for the project based on the effect size obtained in our dose-response study.^[4]^ Regarding the effect of intranasal oxytocin (24 IU at a latency of 45 minutes) on the amygdala response to high intensity fearful faces we observed an effect size of *dz* = 0.56 in a within-subject design. To detect an oxytocin effect of this size (with α = 0.05 and power = 0.75), we needed to test at least 48 participants in a between-subject design (i.e., 24 subjects in each group). In total, 122 healthy women and 124 healthy men were included in the study to control for drop-outs and exclusions. The final sample included 48 subjects (25 women) in the PLC_tra_ and PLC_int_ group, 62 subjects (31 subjects) in the PLC_tra_ and OXT_int_ group, 56 subjects (25 subjects) in the EST_tra_ and PLC_int_ group, and 52 subjects (27 women) in the EST_tra_ and OXT_int_ group.

**Experimental design**

Questionnaires assessing mood (Positive and Negative Affect Schedule [PANAS])^[5]^ and state anxiety (State-Trait Anxiety Inventory [STAI])^[6]^ were administered twice, first directly following the EST_tra_ or PLC_tra_ treatment at the beginning of the testing session and after the fMRI session. The study was carried out at the University Hospital Bonn. The data collection started in October 2018 and was completed in January 2020. Data collection was completed before the start of the COVID-19 pandemic. M.C., I.T. and B.A. enrolled participants and assigned participants to the treatment based on the random allocation sequence (double-blind) generated by D.S.. To generate the random allocation sequence, an integer generator was used (https://www.random.org/).

**Memory performance**

To quantify memory performance, a combined score involving the number of correct responses (hits) and number of false alarms is often used, since neither one should be interpreted without the other.^[7]^ Simply calculating hits minus false alarms would lead to heavily skewed scores in the case of many false alarms. Thus, both scores were z-standardized allowing for comparisons of measures with different ranges of absolute values.^[8]^

**Statistical analyses**

Demographic and psychological data (i.e., age, autistic-like traits, depressive symptoms, alexithymia, trait anxiety and social anxiety) were considered covariates in the main analyses with significant behavioural (*d’*) and neural outcomes (i.e., parameter estimates of significant contrasts of interests). As an additional explorative measure, confidence ratings were used to calculate *Meta d’* for each participant, which reflects the participants’ metacognitive sensitivity and thus the efficacy in which the confidence ratings of the participants discriminate between correct and incorrect judgements. *Meta d’* can be directly compared to *d’*. If *meta d’* equaled *d’*, the participant was acting in a metacognitively ideal manner. If *Meta d’* was not equal to *d’*, the participants either underperformed or outperformed the expectation.^[9,10]^ To compute *Meta d’*, we used code provided by Maniscalco and Lau (<http://www.columbia.edu/~bsm2105/type2sdt/>).^[9]^

**fMRI data acquisition and analysis**

Functional data were obtained using a T2*-weighted echoplanar (EPI) sequence [repetition time (TR) = 2690 ms, echo time (TE) = 30 ms, ascending slicing, matrix size: 96 x 96, voxel size: 2 x 2 x 3 mm³, slice thickness = 3.0 mm, distance factor = 10%, field of view (FoV) = 192 mm, flip angle = 90°, 41 axial slices]. High-resolution T1-weighted structural images were collected on the same scanner (TR = 1660 ms, TE = 2.54 ms, matrix size: 256 x 256, voxel size: 0.8 x 0.8 x 0.8 mm³, slice thickness = 0.8 mm, FoV = 256 mm, flip angle = 9°, 208 sagittal slices). To control for inhomogeneity of the magnetic field, fieldmaps were obtained for each T2*-weighted EPI sequence and included during preprocessing of the fMRI data (TR = 392 ms, TE [1] = 4.92, TE [2] = 7.38, matrix size: 64 x 64, voxel size: 3 x 3 x 3, slice thickness = 3.0 mm, distance factor = 10%, FoV = 192 mm, flip angle 60°, 37 axial slices).

*Preprocessing*

The first five volumes of each functional time series were discarded to allow for T1 equilibration. Functional images were corrected for head movements between scans by an affine registration. Images were initially realigned to the first image of the time series before being re-realigned to the mean of all images. To correct for signal distortion based on B0-field inhomogeneity, the images were unwarped by applying the voxel displacement map (VDM file) to the EPI time series (Realign & Unwarp). Normalization parameters were determined by segmentation and nonlinear warping of the structural scan to reference tissue probability maps in Montreal Neurological Institute (MNI) space. Normalization parameters were then applied to all functional images, which were resampled at 2 x 2 x 2 mm³ voxel size. For spatial smoothing, a 6-mm full width at half maximum (FWHM) Gaussian kernel was used. Raw time series were detrended using a high-pass filter (cut-off period 128 s).

*fMRI data analyses*

In addition to the model described in the main text, we used another model with twelve conditions (valence (3) x memory (2) x sociality (2)) which were modelled by a stick function convolved with a haemodynamic response function. Button presses were included as regressors of no interest. Furthermore, to exclude scans of participants with excessive head movements, the artefact detection toolbox was used (ART; http://www.nitrc.org/projects/artifact_detect) to identify high motion volumes using a volume-to-volume shift of >1.5 mm and a volume-to-volume change in mean signal intensity of >3 standard deviations. Artefacts were treated as regressors of no interest in the following analysis. Subjects with >20% volumes identified as outliers by ART were excluded (n = 1 participant had to be excluded, but this person did not complete the surprise recognition task either).

On the first level, task-specific effects were modelled (e.g.: [Remembered > Forgotten], [Negative and Positive _remembered_ > Negative and Positive _forgotten_], [Negative _remembered_ > Negative _forgotten_], [Neutral _remembered_ > Neutral _forgotten_], [Positive _remembered_ > Positive _forgotten_], [Negative and Positive _remembered > forgotten_ > Neutral _remembered > forgotten_], [Negative _remembered > forgotten_ > Neutral _remembered > forgotten_], and [Positive _remembered > forgotten_ > Neutral _remembered > forgotten_]).

To further examine the potential influence of sex, estradiol, and oxytocin on task-based functional connectivity, a generalized psychophysiological interaction (gPPI) analysis was conducted with the anatomical regions-of-interest (ROIs) as seeds. The analysis was operated with the same preprocessed data, regressors, and contrasts that were used in the SPM analyses. The CONN toolbox (v19.c., www.nitrc.org/projects/conn, RRID:SCR_009550)^[11]^ was used to analyse task-based functional connectivity.

**Questionnaires**

To characterize the sample, we assessed depressive symptoms (Becks Depression Inventory [BDI])^[12]^, alexithymia (Toronto Alexithymia Scale [TAS])^[13]^, social anxiety (Liebowitz Social Anxiety Scale [LSAS])^[14]^, autistic-like traits (Autism Spectrum Quotient [AQ])^[15]^ and trait anxiety (State Trait Anxiety Inventory [STAI])^[6]^. For the presentation of the questionnaires, Qualtrics software was used (Provo, USA). To measure baseline memory performance, the participants completed an adapted version of the Verbal Learning and Memory Test (VLMT).^[16]^ In line with our preregistration, we planned to exclude subjects with an abnormal memory (i.e., learning performance ± 3 SD different from the mean). None of the participants showed abnormal memory performance, thus resulting in no further exclusions.

**Neuroendocrine parameters**

*1.8.1. Estradiol and other gonadal hormones*

Serum estradiol, testosterone and progesterone were determined by fully automated electrochemiluminescent immunoassays (ECLIA, Elecsys test) on a cobas e801 analyzer (Roche Diagnostics, Mannheim, Germany) according to the manufacturer’s instructions (Roche Diagnostics). The coefficients of variation for intra-assay and inter-assay precision were 1.63 % and 2.51 % for estradiol, 2.27 % and 3.71 % for testosterone, and 2.28 % and 2.83 % for progesterone, respectively.

*Oxytocin*

Plasma samples for the measurement of oxytocin (OXT) concentrations were collected with commercial sampling devices (Vacuette, Greiner Bio-One International, Austria) containing ethylenediaminetetraacetic acid (EDTA) and aprotinin. Vacuettes were immediately centrifuged at 3250 rpm for 10 minutes, and aliquoted samples were stored at -80°C until assayed. OXT concentrations were extracted and quantified using a highly sensitive and specific radioimmunoassay (RIAgnosis, Munich, Germany). The limit of detection was 0.1-0.5 pg, depending on the age of the tracer. Intra-assay and inter-assay coefficients of variability were < 10%. All samples to be compared were assayed in the same batch, i.e., under intra-assay conditions.

**fMRI picture set**

The stimuli in the fMRI task were selected based on a pilot study. A sample of 36 healthy subjects (16 females) was shown 360 stimuli, divided into six blocks, each containing 60 pictures. The valence of the pictures was positive, neutral, or negative, and the content of the pictures was either social (defined as the presence of a depicted human) or nonsocial. The pictures were selected from the IAPS database^[17]^ and from the internet. The participants were asked to rate the valence and arousal of each stimulus on a 7-point Likert scale. Based on their ratings, the picture set for the fMRI task (n = 120) and 60 distractor pictures for the surprise recognition task were chosen such that negative and positive stimuli produced comparable arousal ratings and sex differences were absent for all ratings.

The final picture set for the fMRI task consisted of 120 pictures, with 20 pictures in each category. Valence (*F*_(2,68)_ = 260.84, *p* < 0.001, η_p_^2^ = 0.89, BF_incl_ = 5.603 x 10^41^) and arousal ratings (*F*_(2,68)_ =51.21, *p* < 0.001, η_p_^2^ = 0.60, BF_incl_ = 2.569 x 10^12^) differed significantly between the three valence categories, but there were no main or interaction effects of sex (*p* > 0.05; all BF_incl_ < 0.4). Post-hoc comparisons of the valence ratings revealed significant differences between all valence categories (negative and neutral: *t*_(35)_ = -13.94, *p* < 0.001, *d* = -2.32, BF_10_ = 8.152 x 10^12^; negative and positive: *t*_(35)_ = -17.37, *p* < 0.001, *d* = 2.90, BF_10_ = 5.430 x 10^15^; positive and neutral: *t*_(35)_ = 16.53, *p* < 0.001, *d* = -2.75, BF_10_ = 1.212 x 10^15^). Further post hoc comparisons revealed that the arousal ratings between negative and positive stimuli were comparable (*t*_(35)_ = 0.09, *p* = 0.93, *d* = 0.01; BF_10_ = 0.18), whereas the neutral valence category was perceived as less arousing than both the positive (*t*_(35)_ = -11.88, *p* < 0.001, *d* = -1.98, BF_10_ = 9.405 x 10^10^) and negative (*t*_(35)_ = 8.18, *p* < 0.001, *d* = 1.36, BF_10_ = 9.263 x 10^6^) valence categories. The 60 distractor pictures in the surprise recognition task, with 10 pictures in each category, were carefully selected to match the arousal and valence ratings of the picture set used in the fMRI task.

**Supplementary Results**

**Behavioural results**

Across sex and treatment groups, we also observed a significant interaction between valence and sociality (*F*_(2,388)_ = 5.89, *p* < 0.01, η_p_^2^ = 0.03) such that the effect of sociality was significantly more pronounced for positive items (*t*_(201)_ = 4.24 , *p* < 0.001, *d* = 0.30; BF_10_ = 388.63), than for neutral (*t*_(201)_ = 1.90 , *p* = 0.06, *d* = 0.13; BF_10_ = 0.46) and negative (*t*_(201)_ = -0.46 , *p* = 0.64, *d* = -0.03; BF_10_ = 0.09) stimuli. Given that the effect of sociality was only evident for positive items, it seems unlikely that the behavioral task induced an attentional bias towards social stimuli *per se*. Furthermore, we repeated the behavioural analyses with a more liberal recognition criterion in which an item was classified as remembered if the participants correctly identified the item in the recognition task irrespective of the confidence of the classification. These analyses yielded a similar pattern of results.

**Meta D’**

We observed a significant three-way interaction of sex * EST_tra_ treatment * OXT_int_ treatment on *Meta d’* (cf. **Table S1**; *F*_(1,189)_ = 5.23, *p* = 0.02, η_p_^2^ = 0.03; BF_incl_ = 2.72). However, there were no significant sex differences in either treatment type (all *p*s_cor_ > 0.05; all BF_10_ < 2.5) and the interaction between the EST_tra_ and OXT_int_ treatments was not significant in either men (*F*_(1,92)_ = 4.99, *p* = 0.03 [*p*_cor_ = 0.06], η_p_*^2^* = 0.05; BF_incl_ = 2.22) or women (*F*_(1,97)_ = 1.55, *p* = 0.22, η_p_*^2^* = 0.02; BF_incl_ = 0.52).

*Difference score D’ and Meta D’*

As *Meta d’* can be directly compared to d’, we calculated a difference score between *meta d’* and *d’* for each participant. Therefore, we calculated a mixed-design ANOVA with EST_tra_ treatment, OXT_int_ treatment, and sex as between-subject factors and the difference score as dependent variable. However, analyses revealed no significant main or interaction effects of sex and treatment type (all *p*s > 0.05; all BF_10_ < 0.7). Further analyses of the male and female subgroups revealed that in each of the four treatment conditions, the difference score was not significantly different from zero (all *ps* > 0.05; all BF_10_ < 0.6). Thus, in conclusion, participants acted metacognitively ideal, and neither sex nor the treatment types significantly changed meta-cognition.

**Whole brain**

Analyses revealed that there were neither significant whole-brain main effects of sex or treatment type (EST_tra_ or OXT_int_ treatment), nor a significant three-way interactions (sex * EST_tra_ treatment * OXT_int_ treatment) for the contrasts of interests (1. [Remembered > Forgotten] and 2. [Emotional _Remembered > Forgotten_ > Neutral _Remembered > Forgotten_]; all *p*s_cor_ > 0.05). For whole brain task effects see Supplementary **Tables S2-3**.

**Neural emotional memory effect**

A significant sex * EST_tra_ treatment * OXT_int_ treatment interaction emerged for the emotional memory effect in the left hippocampus ([Emotional _Remembered > Forgotten_ > Neutral _Remembered > Forgotten_]; MNI peak coordinates [x, y, z]: -14, −40, 10, *F*_(1,194)_ *=* 16.66, on peak level *p*_FWE_ = 0.019).

*Sex differences*

Post hoc tests to unravel the significant sex * EST_tra_ treatment * OXT_int_ treatment interaction for the emotional memory effect in the left hippocampus revealed that women following placebo administration exhibited nonsignificantly larger left hippocampus responses to emotional remembered stimuli than men (i.e., contrast of interest: [Emotional _Remembered > Forgotten_ > Neutral _Remembered > Forgotten_]; PLC_tra_ & PLC_int_; *t*_(42)_ = -2.42, *p* = 0.02 [*p*_cor_ = 0.08], *d* = -0.74; BF_10_ = 2.88). This sex difference was reversed (i.e. male > female) after OXT_int_ treatment (PLC_tra_ & OXT_int_; *t*_(54)_ = 2.69, *p* = 0.01 [*p*_cor_ = 0.04], *d* = 0.72; BF_10_ = 4.91) and EST_tra_ treatment (EST_tra_ & PLC_int_: *t*_(52)_ = 1.92, *p* = 0.06, [*p*_cor_ = 0.24], *d* = 0.52; BF_10_ = 1.23). Interestingly, sex differences were absent in the combined treatment group (*t*_(46)_ = -1.27, *p* = 0.21, [*p*_cor_ = 0.84], *d* = -0.37; BF_10_ = 0.55).

*Treatment effects*

We found a significant interaction between the OXT_int_ and EST_tra_ treatments for the emotional memory effect in the left hippocampus in men ([Emotional _Remembered > Forgotten_ > Neutral _Remembered > Forgotten_]; *F*_(1,95)_ = 9.54, *p* < 0.01 [*p*_cor_ < 0.01]*,* η_p_^2^ *=* 0.09; BF_incl_ = 14.29) and in women (*F*_(1,99)_ = 6.86, *p* = 0.01 [*p*_cor_ = 0.02]*,* η_p_^2^ *=* 0.07; BF_incl_ = 4.70). Thus, given these data, the Bayes factors indicated that models including the interaction between OXT_int_ and EST_tra_ were 14.29 and 4.70 times more likely than models without it. In men, OXT_int_ nonsignificantly increased hippocampal responses after PLC_tra_ (PLC_tra_ & OXT_int_ vs. PLC_tra_ & PLC_int_ *t*_(44)_ = -2.53*, p* = 0.02 [*p*_cor_ = 0.06], *d* = -0.76; BF_10_ = 3.57*)*, but nonsignificantly reduced the response after EST_tra_ treatment (EST_tra_ & OXT_int_ vs. EST_tra_ & PLC_int_ *t*_(51)_ = 1.97*, p* = 0.05 [*p*_cor_ = 0.2], *d* = 0.54; BF_10_ = 1.34). EST_tra_ had no significant effect on hippocampal activation in men after PLC_int_ (EST_tra_ & PLC_int_ vs. PLC_tra_ & PLC_int_: *t*_(46)_ = -1.59*, p* = 0.12 [*p*_cor_ = 0.48], *d* = -0.47; BF_10_ = 0.81), but significantly decreased the hippocampus activation in the combined group (EST_tra_ & OXT_int_ vs. PLC_tra_ & OXT_int_: *t*_(49)_ = 2.89*, p* < 0.01 [*p*_cor_ = 0.02], *d* = 0.81; BF_10_ = 7.49). In women, OXT_int_ nonsignificantly decreased hippocampal responses after PLC_tra_ (PLC_tra_ & OXT_int_ vs. PLC_tra_ & PLC_int_ *t*_(52)_ = 2.54*, p* = 0.01 [*p*_cor_ = 0.06], *d* = 0.69; BF_10_ = 3.67), but had the opposite effect after EST_tra_ treatment (EST_tra_ & OXT_int_ vs. EST_tra_ & PLC_int_ *t*_(47)_ = -1.17*, p* = 0.25 [*p*_cor_ *≈* 1], *d* = -0.34 ; BF_10_ = 0.50). Likewise, EST_tra_ significantly decreased hippocampal activation in women after PLC_int_ (EST_tra_ & PLC_int_ vs. PLC_tra_ & PLC_int_: *t*_(48)_ = 2.84*, p* < 0.01 [*p*_cor_ = 0.03], *d* = 0.80; BF_10_ = 6.74) and had no effects in the combined group (EST_tra_ & OXT_int_ vs. EST_tra_ & PLC_int_ *t*_(51)_ = -0.88*, p* = 0.38 [*p*_cor_ *≈* 1], *d* = -0.24; BF_10_ = 0.38).

**Simple effect analyses of the oxytocin treatment at the neural level**

To examine a possible treatment effect of OXT_int_, two-sample *t*-tests were calculated to compare the neural responses in the PLC_tra_ & PLC_int_ and PLC_tra_ & OXT_int_ groups separately for women and men. There were no significant OXT_int_ treatment effects in women or men for the two main contrasts ([Remembered > Forgotten]) at the whole brain level, or in the amygdala or insula ROIs (all *p*s > 0.05).

**Simple effect analyses of the estradiol treatment at the neural level**

To examine a possible treatment effect of EST_tra_, two-sample *t*-tests were calculated to compare the neural responses in the PLC_tra_ & PLC_int_ and EST_tra_ & PLC_int_ groups separately for women and men. There were no significant EST_tra_ treatment effects in women or men for the contrast ([Remembered > Forgotten) at the whole brain level, or in the amygdala or insula ROIs (all *p*s > 0.05).

**Simple effect analyses of sex at the neural level**

To examine possible sex effects, two-sample *t*-tests were calculated to compare the neural responses of women and men separately for the four treatment groups. In the combined treatment group (EST_tra_ &OXT_int_), men showed an increased activation in the left insula in contrast to women for the memory effect ([Remembered > Forgotten]; MNI peak coordinates [x, y, z]: -32, 18, -4, *t*_(46)_ *=* 4.59, on peak level *p*_FWE_ = 0.013). Further comparisons at the whole brain level and the remaining ROIs were not significant for the Remembered > Forgotten contrast (all *p*s > 0.05).

**Social vs. nonsocial stimuli**

To investigate social-specific neural treatment effects, we calculated a mixed-design ANOVA with EST_tra_ treatment, OXT_int_ treatment, and sex as between-subject factors and the neural responses to social compared to non-social stimuli as dependent variables (i.e., 1. [Social > Nonsocial]; 2. [Social Remembered > Nonsocial Remembered]). We found no significant main or interaction effects of sex and treatment types for the hippocampus, insula and amygdala responses in these contrasts. In conclusion, neither sex nor treatment type significantly changed neural activity as a function of sociality.

**Functional connectivity**

*Three-way interactions*

To address the effects of the two treatment types on task-based functional connectivity, we conducted a generalized psychophysiological interaction (gPPI) analysis using the CONN Toolbox. The hippocampus, insula and amygdala were used as seeds in whole-brain seed-to-voxel analyses. To investigate a potential three-way interaction, the memory-related connectivity values (i.e., contrast of interest: [Remembered > Forgotten] and [Emotional _Remembered > Forgotten_ > Neutral _Remembered > Forgotten_]) were used as dependent variables in a mixed-design ANOVA. The analyses did not reveal significant three-way interactions with sex and treatment types for either contrast.

*Simple effect analyses of the treatments and sex for [Remembered > Forgotten]*

In addition, exploratory analyses and post-hoc *t*-tests revealed no simple effects of treatment or sex (e.g. men vs. women in the placebo group (PLC_tra_ & PLC_int_)) for the contrast [Remembered > Forgotten].

**Supraphysiological EST levels**

Women exhibited a significantly larger increase in blood EST levels than men. To address that difference, we included the EST increase as a covariate in our analyses. To further probe the impact of this difference, we used a median-dichotomization and excluded EST_tra_-treated women with large EST increase (n = 23). In this subsample, the treatment-induced increases in EST levels were comparable between women and men within the treatment groups (all *p*s > 0.05; BF_10_ < 0.6) We repeated the main behavioral and neural analyses with this subsample and observed a similar pattern of results (see **Figure S1**). In line with our reported main findings, we also found a significant three-way interaction of sex, OXT_int_, and EST_tra_ treatment in the left hippocampus responses to remembered stimuli compared to forgotten stimuli (MNI peak coordinates [x, y, z]: -12, −38, 8, *F*_(1,165)_ = 15.01, *p* < 0.01 [*p*_cor_ < 0.01]*,* η_p_^2^ *=* 0.08; BF_incl_ = 208.56), as well as a significant three-way interaction for the recognition memory (*F*_(1,165)_ = 6.24, *p* = 0.01, η_p_*^2^* = 0.04; BF_incl_ = 5.05). Further analyses of the extracted parameter estimates revealed results that were comparable to our reported main findings. In line with our main analyes, there were no significant sex differences after EST_tra_ treatment (EST_tra_ & PLC_int_; recognition memory: *t*_(41)_ = -0.04, *p* = 0.97, *d* = -0.01; BF_10_ = 0.32; hippocampal response: *t*_(41)_ = 1.28, *p* = 0.21, *d* = 0.42; BF_10_ = 0.60). After the combined treatment (EST_tra_ & OXT_int_), similar, non-significant, sex differences as in the placebo group (PLC_tra_ & PLC_int_) were evident for hippocampal responses (women > men; *t*_(29)_ = -2.13*, p* = 0.04 [*p*_cor_ = 0.16], *d* = -0.84; BF_10_ = 1.83) andrecognition memory (*t*_(29)_ = -1.49, *p* = 0.15, *d* = -0.59; BF_10_ = 0.81). Therefore, the significantly larger EST increase in women than men has no major impact on the results.

**Further hormonal assessments**

In addition to a significant main effect of sex (*F*_(1,180)_ = 771.19, *p* < .001, η_p_^2^ = .81; BF_incl_ = 5.65 x 10^64^), we observed a significant interaction between sex, time, and EST_tra_ treatment (*F*_(1.73,311.46)_ = 3.61, *p* = .03, η_p_^2^ = .02; BF_incl_ = 2.93) for testosterone levels. Separate post-hoc *t*-tests for sex and time points revealed that estradiol-treated participants (EST_tra_ & PLC_int_ and EST_tra_ & OXT_int_) showed non-significantly decreased post-treatment testosterone levels compared to the placebo groups (PLC_tra_ & PLC_int_ and PLC_tra_ & OXT_int_*,*: women *t*_(100)_ = -2.39, *p* = 0.02, [*p*_cor_ = 0.11], *d* = 0.47; BF_10_ = 2.55; men *t*_(96)_ = -2.23, *p* = 0.03, [*p*_cor_ = 0.17], *d* = 0.45; BF_10_ = 1.88). We did not observe any main or interaction effects for the progesterone levels.

**Demographic and psychometric baseline characteristics**

Demographics and baseline psychometric assessments of the participants are displayed in **Table S7**. There were no significant differences between treatment groups within sexes (all *p*s_cor_ > 0.05; all BF_10_ < 0.7). Across treatments men were older than women (age*, t*_(200)_ = 2.29, *p* = 0.02, *d* = 0.3; BF_10_ = 1.74). In addition, women reported significantly higher social anxiety (Liebowitz scale, *t*_(195)_ = -3.57 , *p* < 0.001, *d* = -0.51; BF_10_ = 52.37) and increased trait anxiety (STAI Trait*, t*_(186.31)_ = -2.60, *p* = 0.01, *d* = -0.37; BF_10_ = 3.45) than men. Alexithymia, depressive symptoms, and autistic-like traits were not significantly different between the sexes (all *ps* > 0.05; all BF_10_ < 0.9).

Demographic and psychometric variables were included as covariates in the main analyses of behavioural (*d’*) and neural effects ([Remembered > Forgotten] and [Emotional _Remembered > Forgotten_ > Neutral _Remembered > Forgotten_]. All reported effects remained significant and the covariates were not significant with the exception of a significant effect of autistic-like traits (AQ) in the hippocampal response for the contrast [Emotional _Remembered > Forgotten_ > Neutral _Remembered > Forgotten_] (*F*_(1,176)_ = 7.42, *p* < 0.01, η_p_^2^ = 0.04). Increased AQ scores were associated with increased activation (*r* = 0.203, *p* < 0.01; BF_10_ = 5.21). However, the significant three-way interaction in the hippocampus remained significant despite the significant AQ covariate (*F*_(1,176)_ = 11.83, *p* = 0.001, η_p_^2^ = 0.06; BF_incl_ = 70.46).

**Weight, body mass index and hormonal levels**

Neither the weight nor the body mass index (BMI) of the participants did significantly correlate with the baseline EST or OXT levels in men or women (all *p*s > 0.05; all BF_10_ < 0.2) or with treatment-induced EST_tra_ increases in either sex (all *p*s > 0.05; BF_10_ < 1.0). Thus, the EST_tra_ treatment did not result in different peripheral levels depending on weight or BMI. While OXT_int_ treatment did not produce different peripheral OXT levels depending on the weight or BMI in men (all *p*s > 0.05; BF_10_ < 0.4), both BMI and weight positively correlated with the OXT increase in OXT_int_-treated women (weight: *r*_(51)_ = 0.29, *p* = 0.04; BF_10_ = 1.39; BMI: *r*_(51)_ = 0.30, *p* = 0.03; BF_10_ = 1.63). Thus, OXT_int_ treatment resulted in higher peripheral OXT levels in women with a higher BMI and increased weight.

**Side effects**

Three days following the treatment, the participants were asked to report the side effects of their treatment. No participant experienced severe side effects, but 10.3% of the participants reported side effects consisting of light headache, tiredness, dizziness, sleep disturbance, hilarity, lack of concentration, or circulatory problems. Importantly, the proportion of participants who reported side effects did not significantly differ between the treatment groups (PLC_tra_ & PLC_int_: 16.3%, PLC_tra_ & OXT_int_: 7.5%, EST_tra_ & PLC_int_: 9.6%, EST_tra_ & OXT_int_: 8.5%; *χ*^2^_(3)_ = 2.30, *p* = .51).

**Blinding of treatment**

Both the participants and the experimenters who conducted the experiment were blinded to treatments. Following the MRI scan, the participants were asked to guess which treatment they received. Out of the 104 participants (99 with available estimates) in the OXT groups (PLC_tra_ & OXT_int_ and EST_tra_ & OXT_int_ ), 23 (23.2%; 9 men) believed that they had received OXT, while 23 (24.7%; 11 men) subjects in the placebo groups (PLC_tra_ & PLC_int_ and EST_tra_ & PLC_int_; *n* = 98, 93 with available estimates) believed that they had received verum treatment. Out of the 102 participants in the EST groups (EST_tra_ & PLC_int_ and EST_tra_ & OXT_int_, 100 with available estimates), 22 (22.0%; 12 men) estimated that they had received verum treatment, while 17 (18.5%; 2 men) subjects in the placebo groups (PLC_tra_ & PLC_int_ and PLC_tra_ & OXT_int;_ *n* = 100, 92 with available estimates) assumed that they had received EST. In general, the participants who believed in an OXT treatment also guessed an EST treatment (*r*_(190)_ = 0.39, *p* < 0.001; all BF_10_ = 397232.22). However, the treatment estimates did not significantly correlate with the actual treatments (all *p*s > 0.05; all BF_10_ < 0.2).

**Measurements of mood**

A main effect of time was found for the positive (*F*_(1,181)_ = 72.35, *p* < 0.001, η_p_^2^ = .29; BF_incl_ = 1.943 x 10^12^) and negative (*F*_(1,181)_ = 8.80, *p* < 0.01, η_p_^2^ = .05; BF_incl_ = 4.23) affect measured with the PANAS (the Positive and Negative Affect Schedule) for both sexes (cf. **Table S6**). Positive and negative mood significantly decreased from the beginning of the testing session (mean ± SD positive: 28.69 ± 5.91; negative: 12.15 ± 3.19) to the end (mean ± SD positive: 25.15 ± 6.24; negative: 11.55 ± 2.23), suggesting an increasing fatigue over the time course of the experiment. No further significant main or interaction effects of time and EST_tra_ and OXT_int_ treatments were found for positive or negative mood.

**Confidence ratings**

Confidence ratings were significantly higher for correct than incorrect responses (fMRI stimuli: *t*_(201)_ = 11.11, *p* < 0.001, *d* = 0.78; BF_10_ = 3.566 x 10^19^; distractors: *t*_(197)_ = 35.51, *p* < 0.001, *d* = 2.52; BF_10_ = 6.807 x 10^83^). A mixed-design ANOVA with stimulus type (fMRI remembered, fMRI forgotten, distractor correct rejection, distractor false alarm) yielded a significant main effect of sex (*F*_(1,190)_ = 6.50, *p* = 0.01, η_p_^2^ = .03; BF_incl_ = 0.66) and a significant interaction of sex * stimulus type (*F*_(1.88,358.67)_ = 3.99, *p* = 0.02, η_p_^2^ = 0.02; BF_incl_ = 1.08), with women reporting higher confidence than men for remembered fMRI items and correctly rejected distractors, but there were no further main or interaction effects of sex and treatment groups on the confidence ratings (all *p*s > 0.05; all BF_10_ < 0.3).

**Missing values**

The following blood samples were missing due to problems in sample assessment or analysis: baseline (oxytocin, n = 4; estradiol, n = 4; progesterone, n = 5; testosterone, n = 4), post-treatment (estradiol, n = 2; progesterone, n = 1; testosterone, n = 2), and three days after the treatment (estradiol, n = 8; progesterone, n = 8; testosterone, n = 9). Connection issues and technical errors resulted in the loss of questionnaires evaluating depressive symptoms (n = 8), autistic-like traits (n = 4), alexithymia (n = 4), trait anxiety (n = 5), social anxiety (n = 5), treatment guesses (n = 10), pretreatment negative and positive mood (n = 3), and posttreatment negative and positive mood (n = 10). In addition, five *Meta d’* values were missing because five subjects did not have any false alarms in the memory recognition task.

**Attentional control**

The results of the attention control in the fMRI task showed a high rate of correct responses (mean percent ± SD correct responses: 96.90% ± 8.80) which were not influenced by treatment type and sex (all *p*s > 0.05). In combination with a high response count (mean percent ± SD total responses: 98.76 ± 5.36), this demonstrated that the participants were paying attention and understood the task.

estradiol has opposing effects on fairness framing in women and men. *European*

*Neuropsychopharmacology*, *50*, 46-54.

DOI: https://doi.org/10.1016/j.euroneuro.2021.04.006

[2]	Eisenegger, C., von Eckardstein, A., Fehr, E., & von Eckardstein, S. (2013). Pharmacokinetics

of testosterone and estradiol gel preparations in healthy young men. *Psychoneuroendocrinology*, *38*(2), 171-178.

DOI: https://doi.org/10.1016/j.psyneuen.2012.05.018

[3]	Faul, F., Erdfelder, E., Lang, A. G., & Buchner, A. (2007). G* Power 3: A flexible statistical power

analysis program for the social, behavioral, and biomedical sciences. *Behavior research methods*, *39*(2), 175-191. DOI: https://doi.org/10.3758/BF03193146

[4]	Spengler, F. B., Scheele, D., Marsh, N., Kofferath, C., Flach, A., Schwarz, S., Stoffel-Wagner,

B., Maier, W. & Hurlemann, R. (2017). Oxytocin facilitates reciprocity in social communication.

*Social Cognitive and Affective Neuroscience*, *12*(8), 1325-1333.

DOI: https://doi.org/10.1093/scan/nsx061

[5]	Watson, D., Clark, L. A., & Tellegen, A. (1988). Development and validation of brief measures

of positive and negative affect: the PANAS scales. *Journal of personality and social psychology*,

*54*(6), 1063. DOI: https://doi.org/10.1037/0022-3514.54.6.1063

[6]	Spielberger, C. D. (1970). Manual for the State-Trait Anxietry, Inventory. *Consulting*

*Psychologist*. DOI: https://doi.org/10.1007/978-94-007-0753-5_2825

[7]	Haatveit, B. C., Sundet, K., Hugdahl, K., Ueland, T., Melle, I., & Andreassen, O. A. (2010). The validity of d prime as a working memory index: results from the “Bergen n-back task”. *Journal of Clinical and Experimental Neuropsychology, 32*(8), 871-880.

DOI: https://doi.org/10.1080/13803391003596421

[8]	Macmillan, N. A., & Creelman, C. D. (1990). Response bias: Characteristics of detection theory, threshold theory, and" nonparametric" indexes. *Psychological bulletin, 107*(3), 401.

DOI: https://doi.org/10.1037/0033-2909.107.3.401

[9]	Maniscalco, B., & Lau, H. (2012). A signal detection theoretic approach for estimating metacognitive sensitivity from confidence ratings. *Consciousness and cognition, 21*(1), 422-430. DOI: https://doi.org/10.1016/j.concog.2011.09.021

[10]	Rouault, M., Seow, T., Gillan, C. M., & Fleming, S. M. (2018). Psychiatric symptom dimensions are associated with dissociable shifts in metacognition but not task performance. *Biological psychiatry, 84*(6), 443-451. DOI: https://doi.org/10.1016/j.biopsych.2017.12.017

[11]	Whitfield-Gabrieli, S., & Nieto-Castanon, A. (2012). Conn: a functional connectivity toolbox for correlated and anticorrelated brain networks. *Brain connectivity, 2*(3), 125-141.

DOI: https://doi.org/10.1089/brain.2012.0073

[12]	Beck, A. T., Steer, R. A., & Brown, G. K. (1996). Beck depression inventory.

DOI: https://doi.org/10.1007/978-1-4419-1005-9_441

[13]	Taylor, G. J., Ryan, D., & Bagby, M. (1985). Toward the development of a new self-report alexithymia scale. *Psychotherapy and psychosomatics, 44*(4), 191-199.

DOI: https://doi.org/10.1159/000287912

[14]	Liebowitz, M. R. (1987). Social phobia. *Modern problems of pharmacopsychiatry*.

DOI: https://doi.org/10.1159/000414022

[15]	Baron-Cohen, S., Wheelwright, S., Skinner, R., Martin, J., & Clubley, E. (2001). The autism-spectrum quotient (AQ): Evidence from asperger syndrome/high-functioning autism, malesand females, scientists and mathematicians. *Journal of autism and developmental disorders, 31*(1), 5-17. DOI: https://doi.org/10.1023/a:1005653411471

[16]	Helmstaedter, C., & Durwen, H. (1990). VLMT: Verbaler Lern-und Merkfähigkeitstest: Ein praktikables und differenziertes Instrumentarium zur Prüfung der verbalen Gedächtnisleistungen. *Schweizer Archiv für Neurologie, Neurochirurgie und Psychiatrie*.

DOI: https://doi.org/10.1026//0012-1924.45.4.205

[17]	Lang, P. J. (2005). International affective picture system (IAPS): Affective ratings of pictures and instruction manual. *Technical report*.

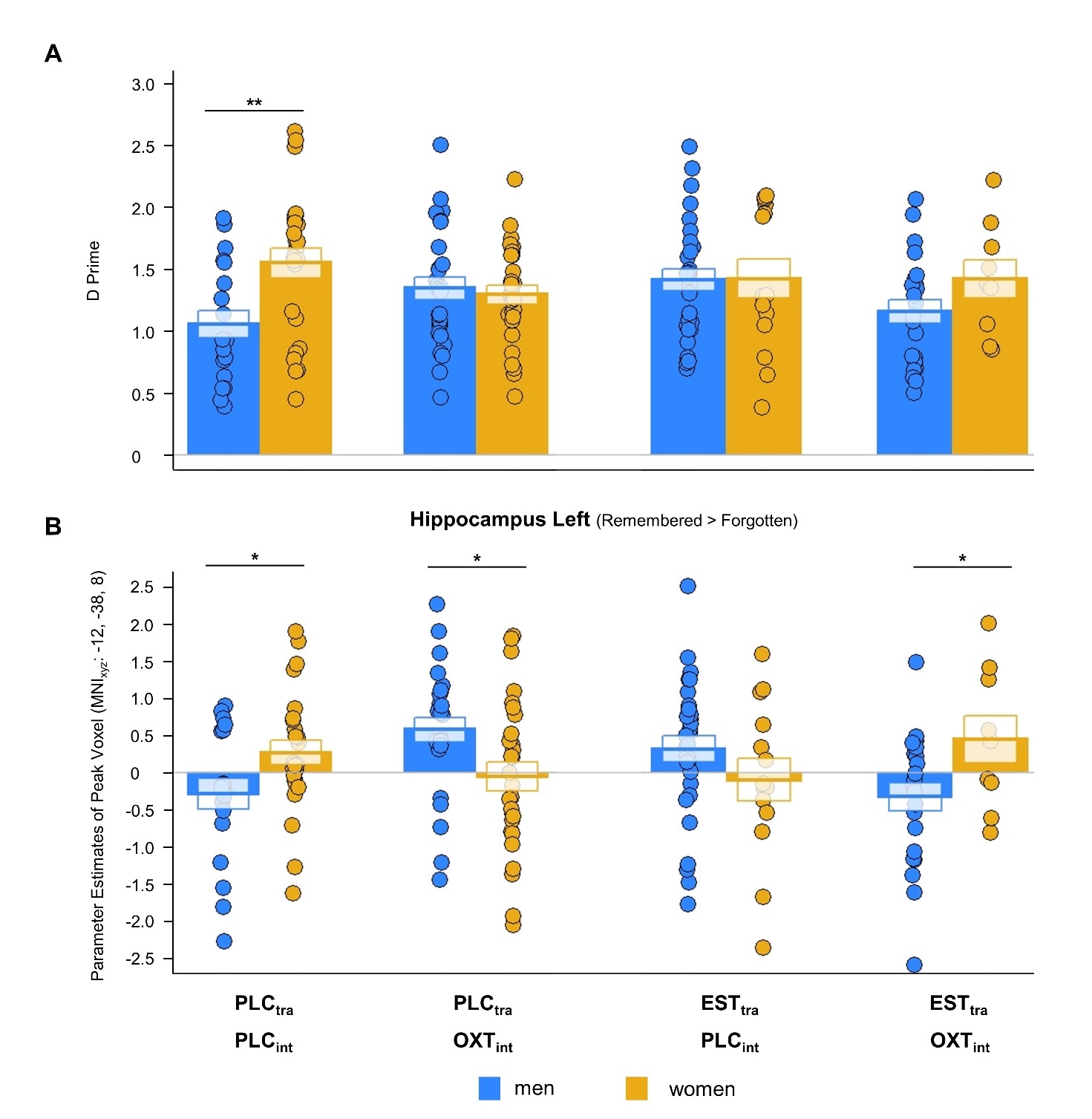
**Supplementary Figures**

**Figure S1.** Treatment effects on mnemonic and hippocampal sex differences in a subsample with comparable blood estradiol (EST) increase in women and men. **A** In line with the results of the total sample, there were no significant sex differences after the single trandermal estradiol treatment (EST_tra_)**.** After the combined treatment, a non-significant sex difference similar to that observed in the placebo group was evident. **B** Further analyses of the parameter estimates for the left hippocampal responses revealed a similar pattern as in the total sample. Intriguingly, the same pattern as that observed in the placebo group was again evident in the combined treatment group. PLC_tra_ = transdermal placebo gel; PLC_int_ = intranasal placebo; OXT_int_ = intranasal oxytocin; EST_tra_ = transdermal estradiol. **p* < 0.05, ***p* < 0.01.

**Supplementary Tables**

**Table S1.** Hit rate, false alarm rate, *D’* and *Meta D’* values in the memory recognition task

|  | Females | | | | Males | | | |
| --- | --- | --- | --- | --- | --- | --- | --- | --- |
|  | PLC_tra_ & PLC_int_  Mean  (n, ± SD) | PLC_tra_ & OXT_int_  Mean  (n, ± SD) | EST_tra_ & PLC_int_  Mean  (n, ± SD) | EST_tra_ & OXT_int_  Mean  (n, ± SD) | PLC_tra_ & PLC_int_  Mean  (n, ± SD) | PLC_tra_ & OXT_int_  Mean  (n, ± SD) | EST_tra_ & PLC_int_  Mean  (n, ± SD) | EST_tra_ & OXT_int_  Mean  (n, ± SD) |
| Hit rate (%) | 0.62  (25, 0.12) | 0.56  (29, 0.15) | 0.63  (25, 0.15) | 0.55  (24, 0.13) | 0.56  (19, 0.17) | 0.57  (27, 0.15) | 0.57  (29, 0.12) | 0.51  (24, 0.21) |
| False alarm rate (%) | 0.14  (25, 0.11) | 0.13  (29, 0.08) | 0.15  (25, 0.10) | 0.12  (24, 0.08) | 0.20  (19, 0.08) | 0.15  (27, 0.09) | 0.13  (29, 0.09) | 0.15  (24, 0.11) |
| *D‘* | 1.55  (25, 0.60) | 1.32  (29, 0.41) | 1.46  (25, 0.52) | 1.42  (24, 0.49) | 1.05  (19, 0.48) | 1.34  (27, 0.48) | 1.41  (29, 0.48) | 1.16  (24, 0.47) |
| *Meta D‘* | 1.55  (24, 0.73) | 1.21  (29, 1.51) | 1.36  (25, 1.50) | 1.65  (23, 1.07) | 1.06  (19, 0.62) | 1.29  (26, 0.98) | 1.55  (29, 0.87) | 0.96  (22, 0.94) |

*Notes.* PLC_tra_ = transdermal placebo gel; PLC_int_ = intranasal placebo; OXT_int_ = intranasal oxytocin; EST_tra_ = transdermal estradiol.

**Table S2.** Whole brain activation for remembered items (i.e., [Remembered > Forgotten]) across sexes and treatments

| Region | Right/Left | Cluster size (voxels) | Peak *F*-score | MNI coordinates | | |
| --- | --- | --- | --- | --- | --- | --- |
|  |  |  |  | x | y | z |
| Inferior temporal gyrus | L | 4889 | 236.45 | -44 | -56 | -10 |
| Middle occipital gyrus | R | 5161 | 184.97 | 36 | -74 | 26 |
| Middle frontal gyrus | R | 4676 | 180.07 | 36 | 52 | 6 |
| Precuneus | R | 3862 | 174.27 | 8 | -68 | 40 |
| Inferior parietal gyrus | R | 1382 | 171.08 | 52 | -48 | 50 |
| Middle frontal gyrus | L | 1705 | 148.42 | -32 | 48 | 8 |
| Inferior frontal gyrus, orbital part | L | 1209 | 133.09 | -36 | 30 | -14 |
| Inferior frontal gyrus | R | 761 | 121.10 | 48 | 32 | 12 |
| Inferior frontal gyrus, orbital part | R | 239 | 105.94 | 26 | 28 | -12 |
| Hippocampus | R | 230 | 105.12 | 20 | -6 | -14 |
| Amygdala | L | 186 | 86.47 | -20 | -8 | -16 |
| Gyrus rectus | L | 203 | 77.99 | -2 | 22 | -18 |
| Precuneus | R | 166 | 74.54 | 16 | -48 | 6 |
| Inferior parietal gyrus | L | 611 | 68.12 | -52 | -54 | 42 |
| Middle temporal gyrus | R | 222 | 64.28 | 66 | -28 | -14 |
| Calcarine fissure | L | 165 | 64.25 | -12 | -48 | 6 |
| Superior frontal gyrus, medial | L | 67 | 49.39 | -6 | 52 | 32 |
| Superior temporal gyrus | R | 36 | 42.32 | 52 | -8 | -14 |
| Superior frontal gyrus | L | 16 | 39.53 | -18 | 10 | 62 |
| Middle frontal gyrus | R | 9 | 33.77 | 50 | -4 | 52 |
| Thalamus | L | 6 | 32.70 | -22 | -26 | 6 |
| Superior temporal gyrus | L | 38 | 31.96 | -58 | -26 | 12 |
| Superior temporal gyrus | R | 30 | 30.92 | 62 | -20 | 10 |
| Insula | R | 6 | 30.34 | 34 | 18 | -10 |
| Hippocampus | L | 3 | 28.81 | -32 | -18 | -14 |
| Median cingulate and paracingulate gyrus | R | 2 | 27.49 | 6 | 2 | 30 |
| Middle temporal gyrus | L | 1 | 26.77 | -54 | -32 | -14 |
| Median cingulate and paracingulate gyrus | L | 1 | 26.04 | -14 | -50 | 36 |

*Notes.* Only clusters with FWE-corrected *p*s < 0.05 on peak level are listed (cluster-forming threshold *p*_FWE_ < 0.05).

**Table S3.** Whole brain activation for the emotional memory effect (i.e., [Emotional _Remembered > Forgotten_ > Neutral _Remembered > Forgotten_]) across sexes and treatments

| Region | Right/Left | Cluster size (voxels) | Peak *F*-score | MNI coordinates | | |
| --- | --- | --- | --- | --- | --- | --- |
|  |  |  |  | x | y | z |
| Occipital lobe | L | 69 | 42.64 | -40 | -56 | -8 |
| Inferior frontal gyrus, triangular part | R | 48 | 40.23 | 48 | 34 | 6 |
| Fusiform gyrus | R | 14 | 34.93 | 42 | -46 | -14 |
| Inferior frontal gyrus, orbital part | L | 10 | 29.54 | -38 | 30 | -14 |
| Middle temporal gyrus | R | 2 | 26.21 | 40 | -62 | 18 |

*Notes.* Only clusters with FWE-corrected *p*s < 0.05 on peak level are listed (cluster-forming threshold *p*_FWE_ < 0.05).

**Table S4.** Estradiol, progesterone and testosterone concentrations at baseline, immediately post treatment and three days after the treatment

|  |  | Females | | | | Males | | | |
| --- | --- | --- | --- | --- | --- | --- | --- | --- | --- |
|  |  | PLC_tra_ & PLC_int_  Mean  (n, ± SD) | PLC_tra_ & OXT_int_  Mean  (n, ± SD) | EST_tra_ & PLC_int_  Mean  (n, ± SD) | EST_tra_ & OXT_int_  Mean  (n, ± SD) | PLC_tra_ &  PLC_int_  Mean  (n, ± SD) | PLC_tra_ & OXT_int_  Mean  (n, ± SD) | EST_tra_ &  PLC_int_  Mean  (n, ± SD) | EST_tra_ &  OXT_int_  Mean  (n, ± SD) |
| Estradiol  (pg/ml) | pre | 34.12  (25, 18.97) | 49.02  (28, 47.04) | 46.60  (24, 25.62) | 44.12  (23, 38.67) | 24.27  (19, 8.04) | 24.94  (27, 11.13) | 22.02  (29, 11.78) | 20.35  (23, 9.77) |
|  | post | 39.12  (25, 20.39) | 50.70  (29, 42.15) | 982.36  (25, 598.66) | 965.09  (23, 381.80) | 27.34  (19, 9.28) | 30.30  (27, 10.82) | 542.93  (29, 439.07) | 623.43  (23, 363.60) |
|  | 3 days post | 45.34  (24, 28.76) | 59.10  (28, 32.74) | 77.66  (23, 58.98) | 58.10  (23, 28.96) | 24.53  (18, 6.47) | 24.12  (26, 8.28) | 23.00  (28, 10.35) | 20.50  (24, 8.11) |
| Progesterone  (ng/ml) | pre | 1.05  (25, 3.54) | 0.71  (28, 2.45) | 0.30  (23, 0.28) | 0.17  (23, 0.09) | 0.20  (19, 0.17) | 0.23  (27, 0.15) | 0.17  (29, 0.11) | 0.13  (23, 0.08) |
|  | post | 1.17  (25, 4.39) | 0.45  (29, 1.48) | 0.19  (25, 0.22) | 0.12  (24, 0.06) | 0.13  (19, 0.10) | 0.13  (27, 0.06) | 0.12  (29, 0.09) | 0.10  (23, 0.05) |
|  | 3 days post | 0.68  (24, 2.58) | 0.23  (28, 0.33) | 0.14  (23, 0.11) | 0.23  (23, 0.44) | 0.16  (18, 0.11) | 0.16  (26, 0.09) | 0.16  (28, 0.13) | 0.14  (24, 0.10) |
| Testosterone  (ng/ml) | pre | 0.28  (25, 0.15) | 0.27  (28, 0.11) | 0.20  (24, 0.14) | 0.24  (23, 0.14) | 4.45  (19, 1.23) | 5.35  (27, 1.85) | 4.70  (29, 1.45) | 4.28  (23, 1.75) |
|  | post | 0.23  (25, 0.13) | 0.25  (29, 0.10) | 0.17  (25, 0.13) | 0.20  (23, 0.12) | 4.67  (19, 1.66) | 5.50  (27, 2.00) | 4.36  (29, 1.69) | 4.38  (23, 1.56) |
|  | 3 days post | 0.27  (24, 0.14) | 0.24  (28, 0.09) | 0.19  (23, 0.14) | 0.24  (23, 0.14) | 4.65  (18, 1.32) | 4.69  (26, 2.11) | 4.71  (28, 1.42) | 4.19  (23, 1.49) |

*Notes.* pre; pretreatment; post, 4.5 hours after the gel administration; 3 days post, 3 days post treatment; PLC_tra_ = transdermal placebo gel; PLC_int_ = intranasal placebo; OXT_int_ = intranasal oxytocin; EST_tra_ = transdermal estradiol.

**Table S5.** Oxytocin baseline and post treatment concentrations

|  | Females | | | | Males | | | |
| --- | --- | --- | --- | --- | --- | --- | --- | --- |
|  | PLC_tra_ & PLC_int_  Mean  (n, ± SD) | PLC_tra_ & OXT_int_  Mean  (n, ± SD) | EST_tra_ & PLC_int_  Mean  (n, ± SD) | EST_tra_ & OXT_int_  Mean  (n, ± SD) | PLC_tra_ & PLC_int_  Mean  (n, ± SD) | PLC_tra_ & OXT_int_  Mean  (n, ± SD) | EST_tra_ & PLC_int_  Mean  (n, ± SD) | EST_tra_ & OXT_int_  Mean  (n, ± SD) |
| Oxytocin pre  (pg/ml) | 1.99  (25, 0.70) | 1.57  (28, 0.66) | 1.66  (24, 0.66) | 1.60  (23, 0.47) | 1.91  (19, 0.82) | 2.01  (27, 0.65) | 1.99  (28, 0.64) | 1.92  (24, 0.67) |
| Oxytocin post  (pg/ml) | 2.11  (25, 0.69) | 5.08  (29, 2.04) | 1.83  (25, 0.61) | 4.79  (24, 1.34) | 2.11  (19, 0.99) | 6.20  (27, 2.78) | 2.27  (29, 0.59) | 6.49  (24, 3.22) |

*Notes.* pre; pretreatment; post, 4.5 hours after the gel administration; PLC_tra_ = transdermal placebo gel; PLC_int_ = intranasal placebo; OXT_int_ = intranasal oxytocin; EST_tra_ = transdermal estradiol.

**Table S6.** Mood measurements

|  | Females | | | | Males | | | |
| --- | --- | --- | --- | --- | --- | --- | --- | --- |
|  | PLC_tra_ & PLC_int_  Mean  (n, ± SD) | PLC_tra_ & OXT_int_  Mean  (n, ± SD) | EST_tra_ & PLC_int_  Mean  (n, ± SD) | EST_tra_ & OXT_int_  Mean  (n, ± SD) | PLC_tra_ & PLC_int_  Mean  (n, ± SD) | PLC_tra_ & OXT_int_  Mean  (n, ± SD) | EST_tra_ & PLC_int_  Mean  (n, ± SD) | EST_tra_ & OXT_int_  Mean  (n, ± SD) |
| Positive affect pre | 27.46  (24, 6.77) | 28.00  (28, 6.45) | 29.28  (25, 5.70) | 27.63  (24, 5.30) | 30.28  (18, 5.17) | 29.33  (27, 6.65) | 27.83  (29, 5.31) | 30.33  (24, 5.48) |
| Positive affect post | 23.74  (23, 6.98) | 24.81  (27, 6.21) | 23.71  (24, 5.47) | 25.00  (24, 5.98) | 27.53  (17, 5.82) | 24.68  (25, 6.43) | 25.52  (29, 6.23) | 26.91  (23, 6.58) |
| Negative affect pre | 13.50  (24, 5.13) | 11.57  (28, 1.55) | 12.64  (25, 4.80) | 11.71  (24, 1.78) | 12.56  (18, 2.41) | 11.89  (27, 3.31) | 11.62  (29, 1.95) | 12.00  (24, 2.52) |
| Negative affect post | 12.48  (23, 2.17) | 11.00  (27, 1.86) | 11.71  (24, 2.26) | 12.13  (24, 2.31) | 10.94  (17, 1.39) | 11.48  (25, 2.33) | 11.14  (29, 1.85) | 11.52  (23, 3.13) |

*Notes.* Mood was assessed with the Positive and Negative Affect Schedule (PANAS). Abbreviations: pre; pretreatment; post, 4.5 hours after the gel administration; PLC_tra_ = transdermal placebo gel; PLC_int_ = intranasal placebo; OXT_int_ = intranasal oxytocin; EST_tra_ = transdermal estradiol.

**Table S7.** Demographic and psychometric baseline characteristics

|  | Females | | | | Males | | | |
| --- | --- | --- | --- | --- | --- | --- | --- | --- |
|  | PLC_tra_ & PLC_int_  Mean  (n, ± SD) | PLC_tra_ & OXT_int_  Mean  (n, ± SD) | EST_tra_ & PLC_int_  Mean  (n, ± SD) | EST_tra_ & OXT_int_  Mean  (n, ± SD) | PLC_tra_ & PLC_int_  Mean  (n, ± SD) | PLC_tra_ & OXT_int_  Mean  (n, ± SD) | EST_tra_ & PLC_int_  Mean  (n, ± SD) | EST_tra_ & OXT_int_  Mean  (n, ± SD) |
| Age  (years) | 23.68  (25, 4.40) | 24.10  (29, 4.52) | 24.72  (25, 5.16) | 24.38  (24, 5.02) | 24.63  (19, 4.76) | 25.07  (27, 3.49) | 26.90  (29, 4.10) | 25.71  (24, 5.07) |
| Depressive symptoms (BDI^b^) | 4.17  (24, 3.41) | 2.11  (28, 2.57) | 3.30  (23, 3.72) | 3.00  (24, 3.09) | 2.89  (18, 3.92) | 1.65  (26, 2.42) | 1.97  (29, 2.41) | 2.86  (22, 2.92) |
| Autistic-like traits (AQ^c^) | 17.29  (24, 6.69) | 13.67  (27, 3.21) | 14.64  (25, 5.69) | 16.71  (24, 6.60) | 16.22  (18, 4.90) | 14.26  (27, 5.54) | 14.76  (29, 4.70) | 15.50  (24, 5.79) |
| Alexithymia (TAS^d^) | 45.67  (24, 9.98) | 41.04  (27, 9.98) | 44.08  (25, 12.90) | 43.96  (24, 11.12) | 42.33  (18, 7.66) | 43.74  (27, 9.32) | 42.07  (29, 7.71) | 40.46  (24, 9.32) |
| Trait anxiety (STAI^e^) | 38.88  (24, 8.25) | 35.63  (27, 8.65) | 34.24  (25, 7.38) | 36.08  (24, 9.38) | 33.06  (18, 8.00) | 32.37  (27, 6.58) | 34.66  (29, 6.11) | 33.13  (23, 6.20) |
| Social anxiety (Liebowitz Total^f^) | 22.22  (23, 9.75) | 18.48  (27, 13.08) | 16.88  (25, 11.91) | 19.46  (24, 12.73) | 15.89  (18, 13.77) | 11.70  (27, 11.13) | 13.17  (29, 11.20) | 12.67  (24, 12.04) |

*Notes.* Subjects rated their depressive symptoms with the ^b^ BDI (Becks Depression Inventory, Beck et al., 1996). Autistic-like traits were measured with the ^c^ AQ (Autism Spectrum Quotient, Baron-Cohen et al., 2006). Alexithymia was assessed with the ^d^ TAS (Toronto Alexithymia Scale, Taylor et al., 1985). The ^e^ STAI-Trait (State-Trait-Anxiety Inventory, Spielberger, 1970) was used to assess trait anxiety and the ^f^ Liebowitz questionnaire was used to measure social anxiety. PLC_tra_ = transdermal placebo gel; PLC_int_ = intranasal placebo; OXT_int_ = intranasal oxytocin; EST_tra_ = transdermal estradiol.

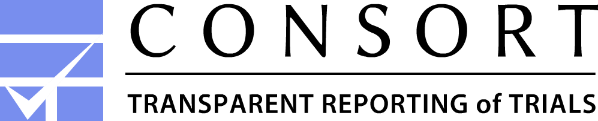
**CONSORT 2010 Flow Diagram**

Dose: 2mg of Placebo Gel

24IU of Placebo Spray

Allocated to intervention (*n* = 55)

♦  Did not receive allocated intervention (*n* = 0)

Males

(*n* = 27)

Excluded from the analysis, due to:

♦ Memory performance (*n* = 7)

♦ Technical issues (*n* = 2)

♦ Anatomical abnormalities (*n* = 1)

♦ Hormonal abnormalities (*n* = 1)

- Analysed (*n* = 44)

Females

(*n* = 28)

Males

(*n* = 19)

Females

(*n* = 25)

Dose: 2mg of Placebo Gel

24IU of Oxytocin Spray

Allocated to intervention (*n* = 71)

♦  Did not receive allocated intervention (*n* = 0)

Males

(*n* = 35)

Excluded from the analysis, due to:

♦ Memory performance (*n* = 8)

♦ Technical issues (*n* = 4)

♦ Discontinuation of study (*n* = 2)

♦ Hormonal abnormalities (*n* = 1)

- Analysed (*n* = 56)

Females

(*n* = 36)

Males

(*n* = 27)

Females

(*n* = 29)

Screening Session (*n* = 295)

And

Excluded (*n* = 49)

♦  did not meet inclusion criteria (*n* = 39)

♦  Discontinued study participation (*n* =10)

### Enrollment

### Allocation to treatment condition

Randomized (*n* = 246)

Dose: 2mg of Estradiol Gel

24IU of Oxytocin Spray

Allocated to intervention (*n* = 56)

♦  Did not receive allocated intervention (*n* = 0)

Males

(*n* = 27)

Excluded from the analysis, due to:

♦ Memory performance (*n* = 4)

♦ Technical issues (*n* = 3)

♦ Anatomical abnormalities (*n* = 1)

- Analysed (n = 48)

Females

(*n* = 29)

Males

(*n* = 24)

Females

(*n* = 24)

Dose: 2mg of Estradiol Gel

24IU of Placebo Spray

Allocated to intervention (*n* = 64)

♦  Did not receive allocated intervention (*n* = 0)

Males

(*n* = 35)

Excluded from the analysis, due to:

♦ Memory performance (*n* = 5)

♦ Technical issues (*n* = 2)

♦ Discontinuation of study (*n* = 1)

♦ Hormonal abnormalities (*n* = 2)

- Analysed (*n* = 54)

Females

(*n* = 29)

Males

(*n* = 29)

Females

(*n* = 25)
